# Loss of the Y chromosome drives epigenetic and transcriptomic plasticity in lung adenocarcinoma

**DOI:** 10.64898/2026.06.02.729627

**Authors:** Kathleen Schlüter, Mei-Ju May Chen, Gizem Altun, Sergio Manzano-Sanchez, Dung-Chi Wu, Nan Zhang, Ofir Griess, Luc Husemann, Fabian Bradic, Joseph Cornick, Nima Esmaeelpour, Oliver Mücke, Riccardo Moro, Katherine Kelly, Sara Chocarro, Etienne Sollier, Albert Fradera-Sola, Claudia Scalera, Maria Jose Alonso-De Gennaro, Siavash Mansouri, Madeleine Dorsch, Patricia Munteanu, Balazs Hegedus, Hauke Winter, Laura V. Klotz, Mark Kriegsmann, Felix J.F. Herth, Thomas Muley, Marc A. Schneider, Daniel Kazdal, Albrecht Stenzinger, Alexander Schramm, Felix J. Hartmann, Pavlo Lutsik, Marcel Schilling, Ursula Klingmüller, Rajkumar Savai, Rocio Sotillo, Barbara M. Grüner, Guy Ron, Efrat Shema, Michael Scherer, Christoph Plass, Maria Llamazares-Prada

## Abstract

Loss of the Y chromosome (LOY) is associated with poor survival across multiple solid tumors, yet the underlying molecular mechanisms remain poorly understood. Here, we identify LOY as a central driver of lineage plasticity and epigenetic heterogeneity in lung adenocarcinoma. Integrating multi-omic profiling of primary samples with isogenic cellular models, we show that LOY triggers epithelial-to-mesenchymal transition (EMT). Mechanistically, LOY causes haploinsufficiency of dosage-sensitive regulators, leading to widespread DNA hypomethylation at EMT gene promoters, including *THY1* and *LOX*. Single-cell multi-omic analyses demonstrate that LOY induces epigenetic heterogeneity, destabilizes the chromatin landscape, and increases lineage plasticity, enabling rapid cellular adaptation to metabolic and genotoxic stress. Moreover, LOY-induced plasticity facilitates tumor engraftment and metastatic dissemination *in vivo*. These findings establish Y-linked gene dosage as a critical guardian of epigenetic stability, providing a mechanistic rationale for how its loss amplifies phenotypic diversity and lineage plasticity, ultimately driving adverse clinical outcomes in LOY patients.

## Introduction

The Y chromosome is essential for male sex determination and contributes to biological differences between males and females in both health and disease^1,2^. Beyond sex-determining loci, the Y chromosome encodes dosage-sensitive genes involved in transcriptional regulation, chromatin remodeling, RNA processing, and protein homeostasis^3^. Subclonal loss of the Y chromosome in blood cells, known as mosaic loss of Y (mLOY), has been associated with impaired immune surveillance, increased all-cause mortality, and elevated susceptibility to Alzheimer’s, chronic kidney disease, cardiovascular disease, and cancer^4–9^.

In addition to aging blood cells, early cytogenetic studies reported loss of the Y chromosome (LOY) in both solid and hematologic malignancies^10–14^. More recently, pan-cancer genome analysis revealed variable LOY prevalence across tumor types and cell compartments, including neoplastic and stromal cells^8,15–18^. LOY is associated with genomic instability, high mutational burden, and *TP53* mutations, altogether contributing to poor clinical outcomes^15,16^. Downstream effects of LOY are context-dependent, influencing tumor progression and therapeutic response^16,19^. In bladder cancer, for instance, LOY reshapes the tumor microenvironment to foster immune evasion while increasing sensitivity to immune checkpoint inhibitors^20^. In uveal melanoma, LOY arises in the absence of *TP53* mutations and is associated with metastasis, underscoring its emerging role in cancer progression^15^. Moreover, LOY is linked to poor prognosis and predicts patient outcomes even in tumors with low LOY frequency, such as prostate adenocarcinoma and mesothelioma^15,16,21,22^.

Non-small cell lung cancer (NSCLC) displays a pronounced sex bias, with men showing higher incidence and mortality than women^23,24^. LOY is prevalent in NSCLC, occurring in 43% of lung adenocarcinoma (LUAD) and 57% of lung squamous cell carcinoma (LUSC) cases in the TCGA cohort^13–16,24,25^. LOY correlates with aggressive disease, increased metastasis, and poor prognosis^24^. Molecularly, LOY tumors show widespread DNA hypomethylation at gene promoters compared to Y-retaining counterparts, potentially driven by loss of *lysine demethylase 5D*, *KDM5D*, and consequent increases in histone 3 lysine 4 tri-methylation (H3K4me3) at promoter regions^24^. Recent studies further suggest that LOY disrupts the regulation of cancer-testis antigens (CTAs) through loss of Y-linked transcription factors and KDM5D, thereby modulating the tumor immune microenvironment^18^. Yet, the mechanisms through which LOY drives tumor aggressiveness and metastasis remain elusive.

Here, we find that LOY induces epigenetic reprogramming by eliminating dosage-sensitive genes on the Y chromosome without compensatory expression from X-linked homologs. We propose that this haploinsufficiency drives tumor cell plasticity and adaptability by amplifying epigenetic heterogeneity. To dissect the functional consequences of LOY in LUAD, we integrated whole-genome and single-cell RNA sequencing data from primary tumors of male LUAD patients, complemented by public datasets. LOY was predominantly detected in malignant cells and, to a lesser extent, within the tumor microenvironment. Using isogenic A549 cell models and primary LUAD samples, we show that LOY activates lineage plasticity through the induction of epithelial-to-mesenchymal transition (EMT) programs. Furthermore, LOY-induced plasticity and epigenetic remodeling confer resilience to metabolic and genotoxic stress *in vitro*. Importantly, this enhanced plasticity translated to a selective advantage *in vivo*, facilitating tumor engraftment and metastasis. Together, these findings establish LOY as a previously unrecognized driver of EMT and tumor plasticity. Ultimately, by linking Y chromosome loss to tumor-cell-intrinsic epigenetic reprogramming, phenotypic diversification, and metastatic capacity, our work provides a mechanistic rationale for the adverse clinical outcomes frequently observed in male LUAD patients with LOY.

## Results

### LOY in LUAD is enriched in tumor cells and associates with adverse clinical outcomes

To characterize LOY in lung adenocarcinoma (LUAD), we performed whole-genome sequencing (WGS) and single-cell RNA sequencing (scRNA-seq) on paired tumor and distal normal lung tissue from four male patients (**Table 1**). We integrated these data with whole-exome sequencing from TCGA (n=237 males) and scRNA-seq from the Lung Cancer Atlas^26^ (LuCA, n=45 males) (**Figure 1A**). WGS revealed specific reductions in Y-chromosome copy number in the tumor regions of two donors (HLD5, presenting subclonal LOY, and HLD25), whereas the Y chromosome was retained in matched distal normal tissues (**Figure 1B**, **Figure S1A**). The modest copy number reduction observed in the distal lung tissue of HLD25 likely reflects mLOY in immune populations, a known risk factor for lung cancer^25,27,28^. Donors HLD21 and HLD35 retained the Y chromosome in both compartments and were classified as ROY tumors (retention of Y chromosome).

**Figure 1:**
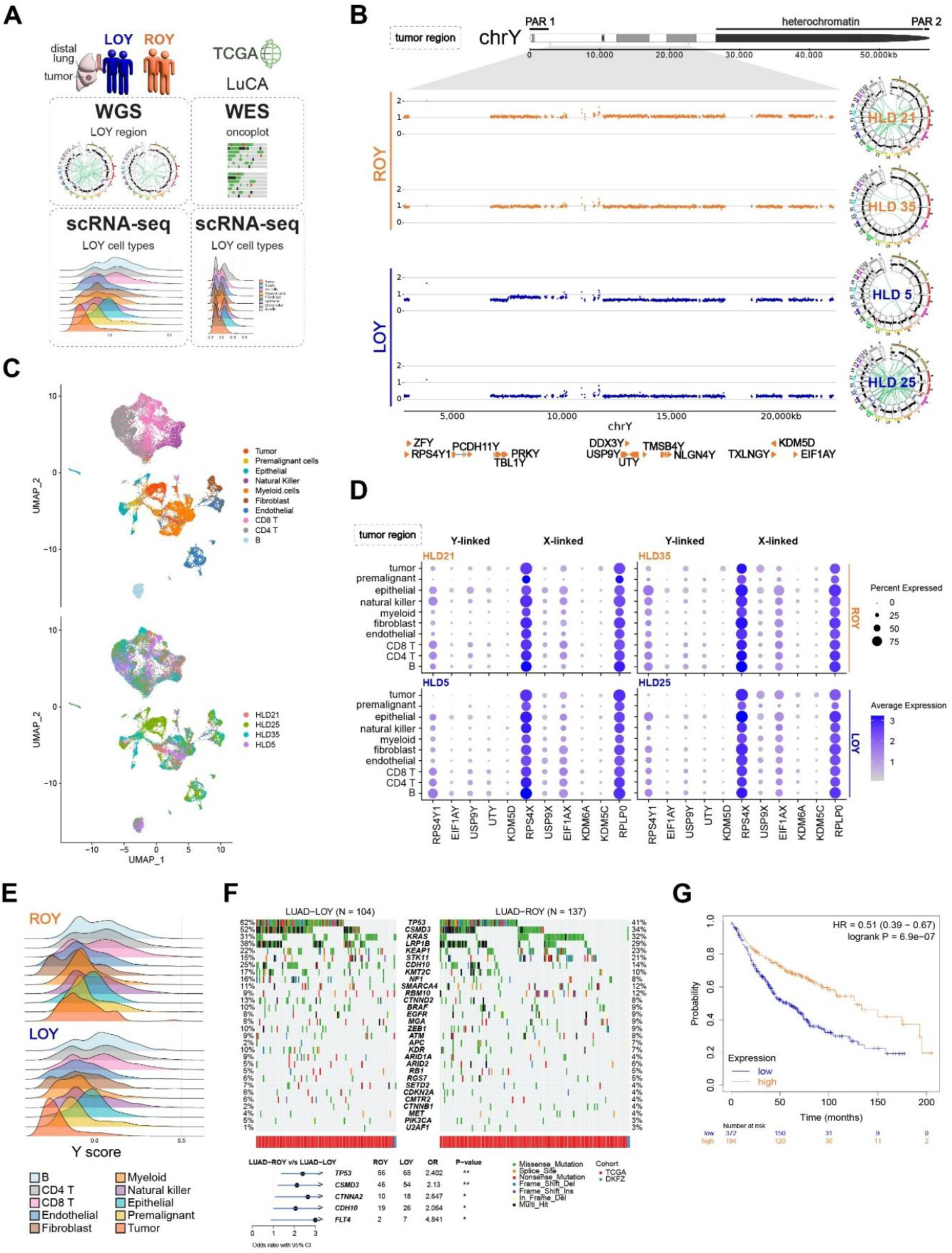
LOY in LUAD is enriched in tumor cells and associates with adverse clinical outcomes. **(A)** Workflow for the characterization of LOY status in lung adenocarcinoma (LUAD) patients from an in-house cohort and public datasets. **(B)** Whole-genome sequencing (WGS) profiles of tumor tissue from two ROY LUAD donors (orange) and two LOY LUAD donors (blue). Zoomed-in views of the Y chromosome (left) and corresponding genome-wide Circos plots (right) are shown. **(C)** UMAP projections of scRNA-seq data from tumor and paired distal lung tissue of ROY (n=2) and LOY (n=2) patients, colored by cell type (top) and patient (bottom). **(D)** Dot plots showing expression levels of five Y-linked genes, their X-linked homologs, and autosomal housekeeping gene *RPLP0* (chromosome 12). Annotated cell types from tumor regions are shown for ROY (top) and LOY (bottom) patient status. Dot size indicates the percentage of cells expressing each gene; color intensity represents mean expression. **(E)** Density plots showing the distribution of Y-scores across all cell types in tumor and paired distal lung samples from ROY (top) and LOY (bottom) patients. **(F)** Oncoplot of the top 30 most frequently mutated genes in TCGA-LUAD and our in-house cohort, stratified by LOY (n = 104, left) and ROY (n = 137, right) status^15^. Fisher’s exact test indicates significant enrichment of mutations in *TP53*, *CSMD3*, *CTNNA2*, *CDH10*, and *FLT4* in LOY tumors (odds ratios shown; OR >1 indicates LOY enrichment. **(G)** Kaplan-Meier survival analysis of 566 male LUAD patients (kmplot.com) stratified by the mean expression of five Y-linked genes (*RPS4Y1*, *EIF1AY*, *USP9Y*, *UTY*, *KDM5D*). Overall survival (OS) is shown for low (blue) and high (orange) expression groups. Median OS: 50 vs. 122 months; HR = 0.51; 95% CI 0.39–0.67, p = 6.9 × 10⁻⁷.

To map LOY to specific cell types, we analyzed scRNA-seq data from tumor (**Figure 1C**) and paired distal lung tissue (**Figure S1B**) across all four patients, spanning ten major cell populations. Expression of five representative Y-linked genes (*RPS4Y1*, *EIF1AY*, *USP9Y*, *UTY*, and *KDM5D*) was absent in tumor cells from LOY donors but preserved in ROY donors and non-malignant cells (**Figure 1D**). Expression of X-linked homologs was maintained across all samples, ruling out technical dropout. We computed a Y-chromosome expression score based on the top 13 Y-linked genes to quantify Y chromosome retention at single-cell resolution^29^. Tumor cells from LOY donors exhibited the lowest Y-scores. In contrast, fibroblasts and myeloid cells displayed bimodal distributions, consistent with mLOY in the tumor microenvironment (TME, **Figure 1E**).

We next linked LOY to malignant states by combining Leiden clustering with inferring copy number changes at the single-cell level. Three out of four malignant epithelial clusters exhibited low Y-scores, while one retained expression (**Figure S1C–E**). Across epithelial cells, the Y-score was inversely correlated with malignancy scores (**Figure S1F**), and LOY clusters displayed elevated stemness and epithelial-to-mesenchymal transition (EMT) scores compared to ROY clusters (**Figure S1G**). These findings were validated in the LuCA dataset^26^ after filtering cells with low gene-detection rates (n = 45, **Figure S1H-I**). Tumor cells exhibited the lowest Y-scores across all cell types. In contrast, immune, epithelial, and mesenchymal cells displayed bimodal Y-score distributions, further pointing towards mLOY in the TME (**Figure S1J**). Using a published random forest classifier to identify LOY at the single-cell level^17^, we confirmed that LOY tumor cells consistently exhibited higher EMT scores than ROY tumor cells, suggesting increased plasticity (**Figure S1L**).

To assess whether LOY tumors exhibit a distinct mutational landscape, we stratified TCGA LUAD samples by Y-status^15^ (102 LOY, 135 ROY tumors). While *TP53* mutations were enriched in LOY tumors, consistent with prior reports^15^, ∼40% of LOY tumors retained wild-type *TP53*, indicating that LOY is not merely a surrogate for *TP53* loss (**Figure 1F**). In our cohort, *TP53* mutations were absent (**Table 2**). In addition, LOY tumors from TCGA showed significant enrichment of *CSMD3*, *CTNNA2*, *CDH10*, and *FLT4* mutations, although the overall mutational landscape was similar between LOY and ROY. To determine the clinical impact of LOY in LUAD, we used the Kaplan–Meier Plotter platform (http://kmplot.com/analysis) and Y-linked gene expression as a proxy for LOY/ROY status. Low Y-linked gene expression was associated with significantly shorter overall survival in male LUAD patients (n = 566; median survival 50 vs. 122 months, **Figure 1G, Figure S1M**). Collectively, these data identify LOY as a frequent event in malignant epithelial cells, associated with aggressive transcriptional states and adverse outcomes.

### LOY drives intrinsic EMT and inflammatory programs

To dissect the direct phenotypic consequences of LOY independent of *TP53* status, we established an isogenic *in vitro* model using the A549 lung adenocarcinoma cell line, which exhibits subclonal LOY^30^. We derived and expanded four independent single-cell LOY and ROY clones (**Figure 2A**), and confirmed complete Y-loss in LOY clones via WGS (**Figure 2B, Figure S2A**).

**Figure 2:**
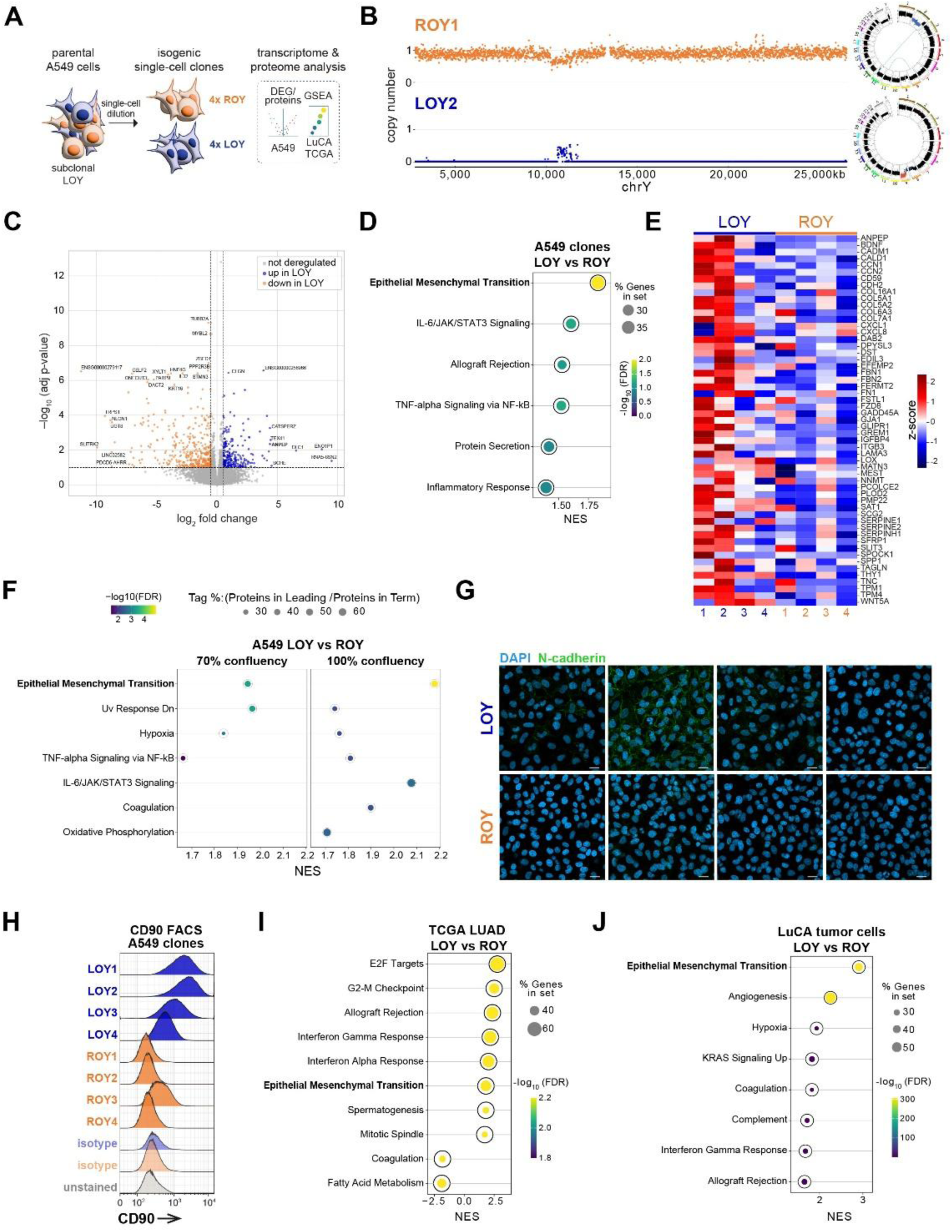
LOY drives intrinsic EMT and inflammatory programs. **(A)** Experimental overview illustrating the generation of isogenic single-cell LOY and ROY clones from parental A549 cells and the multi-omic analysis of clones and patients. **(B)** Representative WGS profiles of ROY (top, orange) and LOY (bottom, blue) clones. Left: zoomed-in view of the Y chromosome. Right: genome-wide Circos plots (right). **(C)** Volcano plot of differentially expressed genes between LOY (n=4) and ROY (n=4) clones (Y-linked genes excluded). X-axis: log2 fold change; y-axis: -log_10_(p-value). Blue: significantly upregulated genes in LOY; orange: downregulated in LOY. **(D)** Dot plot showing the top 6 Hallmark gene sets with FDR<0.25 from preranked GSEA of RNA-seq data. X-axis: normalized enrichment score (NES); y-axis: gene sets. Dot size indicates gene count; color represents -log_10_(FDR). **(E)** Heatmap of lead genes contributing to the EMT gene set enrichment. TPM values are z-scored across clones. **(F)** Dot plot of Hallmark gene sets from GSEA of full proteome data at 70% confluency (left) and 100% confluency (right). X-axis: normalized enrichment score (NES); y-axis: gene sets. Dot size indicates protein count; color represents -log_10_(FDR). (**G**) Confocal microscopy images of clones stained for N-cadherin (green). Nuclei are counterstained with DAPI (4′,6-diamidino-2-phenylindole, blue). Scale bar 20 µm **(H)**. Flow cytometry histograms of CD90 expression for LOY (blue) and ROY (orange) clones, with isotype controls shown. **(I)** Dot plot of top 10 Hallmark gene sets from preranked GSEA of TCGA LUAD samples (LOY vs. ROY). **(J)** Dot plot the top 8 positively enriched Hallmark gene sets from GSEA of the LuCA single-cell dataset^26^ (LOY vs. ROY tumor cells).

Bulk RNA-seq revealed widespread transcriptional rewiring in LOY clones (243 upregulated, 426 downregulated genes; **Figure 2B**, **Table 3**). Consistent with a dosage imbalance hypothesis, we observed no compensatory upregulation of X-linked homologs (**Figure S2C**). Gene set enrichment analysis (GSEA) identified EMT as the top-enriched program in LOY cells, followed by inflammatory response, TNFα, and IL6/JAK/STAT3 signaling, suggesting increased plasticity upon loss of the Y chromosome (**Figure 2D**). While upregulation of EMT genes was consistent, gene expression varied across clones (**Figure 2E**, **Table 3**).

To validate these findings at the protein level, we performed global proteomic profiling of cells grown until 70% or 100% confluency. We observed comparable protein coverage across clones and conditions (**Figure S2C**) and detected Y-chromosome-specific peptides exclusively in ROY cells (**Figure S2D**). Differential expression analysis confirmed a distinct proteome landscape in LOY clones (**Figure S2E**). We identified EMT again as the top-enriched pathway, together with downregulated UV response, hypoxia, and TNFα signaling via NF-κB (**Figure 2F)**. Additionally, LOY cells at high confluency upregulated IL6/JAK/STAT3, coagulation, and oxidative phosphorylation pathways. Moreover, at the single-cell level, LOY clones displayed a variable but evident cadherin switch (increased N-cadherin, decreased E-cadherin), together with increased expression of the stemness marker CD90 (*THY1*) (**Figure 2G-H**, **Figure S2F**).

We found consistent upregulation of EMT and inflammatory pathways in LOY tumors of primary samples from TCGA (bulk) and LuCA^26^ (scRNA-seq) (**Figure 2I-J**), indicating that LOY in tumor cells is associated with activation of EMT, inflammatory signaling, and stemness-associated features.

### Genome-wide epigenetic alterations in A549 LOY cells reshape the epigenetic landscape through DNA hypomethylation

To determine whether the epigenetic remodeling previously observed in patient bulk samples^24^ is driven by tumor-cell intrinsic LOY effects, we profiled CpG methylation and the active promoter mark H3K4me3 using DiMeLo-seq^31,32^ in A549 LOY and ROY clones (**Figure 3A**). Principal component analysis (PCA) separated LOY and ROY clones. LOY clones displayed increased variance along PC2, indicating greater epigenetic heterogeneity (**Figure 3B**). While global DNA methylation was comparable across clones, LOY cells exhibited significant hypomethylation at promoter regions and CpG islands (**Figure S3A**). Analysis of differentially methylated CpG islands revealed prevalent hypomethylation in LOY **(Figure 3C-D)**, underscoring an epigenetic shift characterized by focal hypomethylation at CpG-rich regulatory regions.

**Figure 3:**
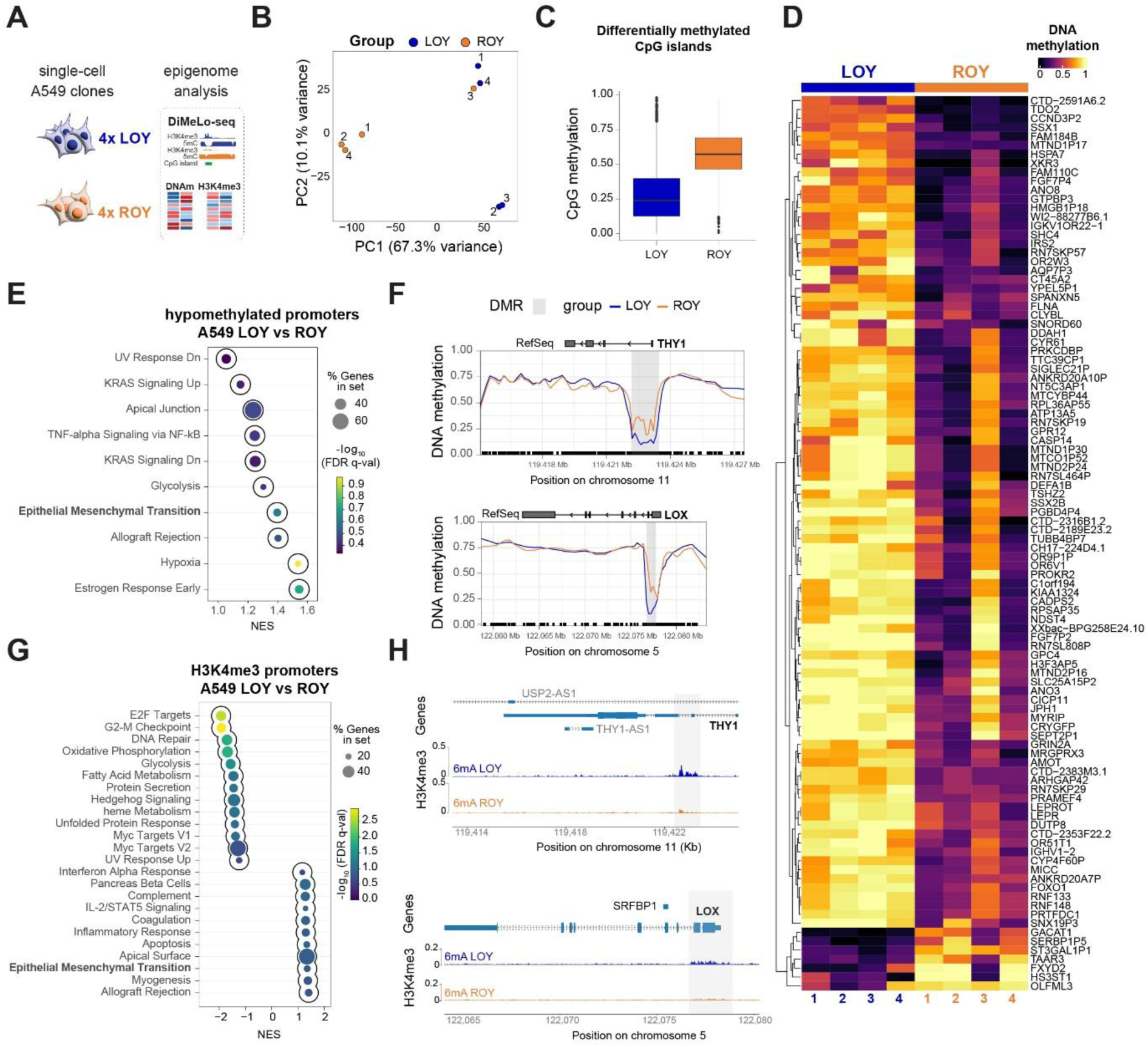
Genome-wide epigenetic alterations in A549 LOY cells reshape the epigenetic landscape through DNA hypomethylation. **(A)** Experimental overview of epigenetic characterization using DiMeLo-seq in isogenic A549 ROY and LOY clones. **(B)** Unsupervised principal component analysis (PCA) of the 50,000 most variable autosomal CpGs. Each dot represents a clone (blue: LOY; orange: ROY). **(C)** Boxplot of differentially methylated CpG islands in LOY (blue) and ROY (orange) cells. **(D)** Heatmap of CpG island methylation levels for autosomal CpG islands within 10 kb of a transcription start site (TSS), filtered to one representative CpG island per gene with the largest absolute methylation difference. Columns represent individual clones, and rows represent associated genes with representative CpG islands. **(E)** GSEA dot plot of promoter-associated DMRs (±5 kb from TSS, Hallmark gene set). Positive normalized enrichment scores (NES) indicate enrichment of genes with hypomethylated promoters in LOY clones. (**F**) DNA methylation tracks at *THY1* (top) and *LOX* (bottom) loci. **(G)** Preranked GSEA of promoter H3K4me3 signal (±2 kb from TSS, MSigDB Hallmark collection). Positive NES indicates increased H3K4me3 in LOY cells. (**H**) Locus-specific profiles of H3K4me3 signal (6mA) averaged for LOY and ROY clones at *THY1* (top) and *LOX* (bottom).

Methylation analysis identified 308,935 differentially methylated CpGs (absolute methylation difference ≥20%), which aggregated into 21,250 differentially methylated regions (DMRs). Notably, 93.0% of these DMRs were hypomethylated in LOY clones **(Figure S3B)**, indicative of a more open and permissive chromatin landscape, consistent with previous data^24^. Most DMRs mapped to intronic and intergenic regions, with a smaller fraction overlapping gene promoters **(Figure S3C)**.

To characterize the regulatory context of these changes, we annotated DMRs to their closest gene (±5 kb around transcription start sites). GSEA of genes with hypomethylated promoters in LOY cells revealed an enrichment for EMT-related gene sets, hypoxia, estrogen response, TNFα signaling, and allograft rejection pathways (**Figure 3E**). We observed focal promoter hypomethylation of the EMT markers *THY1* and *LOX* (*lysyl oxidase*) (**Figure 3F**).

To assess if these DNA methylation changes coincided with altered promoter-associated chromatin activity, we adapted an existing peak-calling approach^33^ to identify H3K4me3 peaks. As expected, H3K4me3-enriched regions exhibited low DNA methylation and localized primarily to promoters (**Figure S3D-F**). We then evaluated differential H3K4me3 enrichment at promoters between LOY and ROY cells. GSEA revealed an enrichment for EMT, allograft rejection, and inflammatory response pathways in LOY cells. Conversely, pathways associated with E2F targets, G2M checkpoint, and DNA repair showed reduced H3K4me3 levels in LOY **(Figure 3G).** Consistent with our methylation data, promoter hypomethylation coincided with increased H3K4me3 at the *THY1* and *LOX* promoters (**Figure 3H**). Furthermore, H3K4me3 signals at EMT gene promoters varied significantly across the clones, as observed in transcriptomic data **(Figure S3G-H**, **Figure 2E)**, highlighting substantial interclonal heterogeneity within the LOY population.

Together, these data indicate that LOY promotes cell plasticity by epigenetic activation at EMT-associated promoters through coordinated DNA hypomethylation and H3K4me3 enrichment, while simultaneously generating significant interclonal chromatin heterogeneity. This complexity suggests that additional histone modifications, such as H3K27ac and H3K27me3, or single-cell analysis are required to fully resolve LOY-associated regulatory states.

### Single-cell profiling reveals LOY-driven epigenetic heterogeneity and transcriptional plasticity

Building on these findings, we hypothesized that haploinsufficiency of Y-linked regulators destabilizes the chromatin landscape, promoting epithelial-to-mesenchymal plasticity and increased single-cell heterogeneity. To test this and to resolve the distinct regulatory states observed in bulk, we profiled isogenic A549 LOY and ROY clones using Epi-CyTOF^34–36^ and single-cell multiome (snRNA-seq/scATAC-seq) (**Figure 4A**). Epi-CyTOF revealed broad remodeling of histone modifications in LOY clones, including enrichment of active marks (H4K16ac, H3K64ac, H3K9ac, H3K27ac, and H3K4me3), repressive marks (H3K9me2), and poised or primed modifications (H3K4me2, H3K27me2/3, and H3K9me1), consistent with global epigenetic remodeling (**Figure 4B**). Interestingly, all the LOY clones showed increased levels of H3K4me3, the target of the Y-linked histone demethylase KDM5D (**Figure S4A-B**). Moreover, LOY clones exhibited higher cell-to-cell variability across histone marks compared with ROY controls, indicating increased histone-state diversity and epigenetic heterogeneity following LOY (**Figure 4C**).

**Figure 4:**
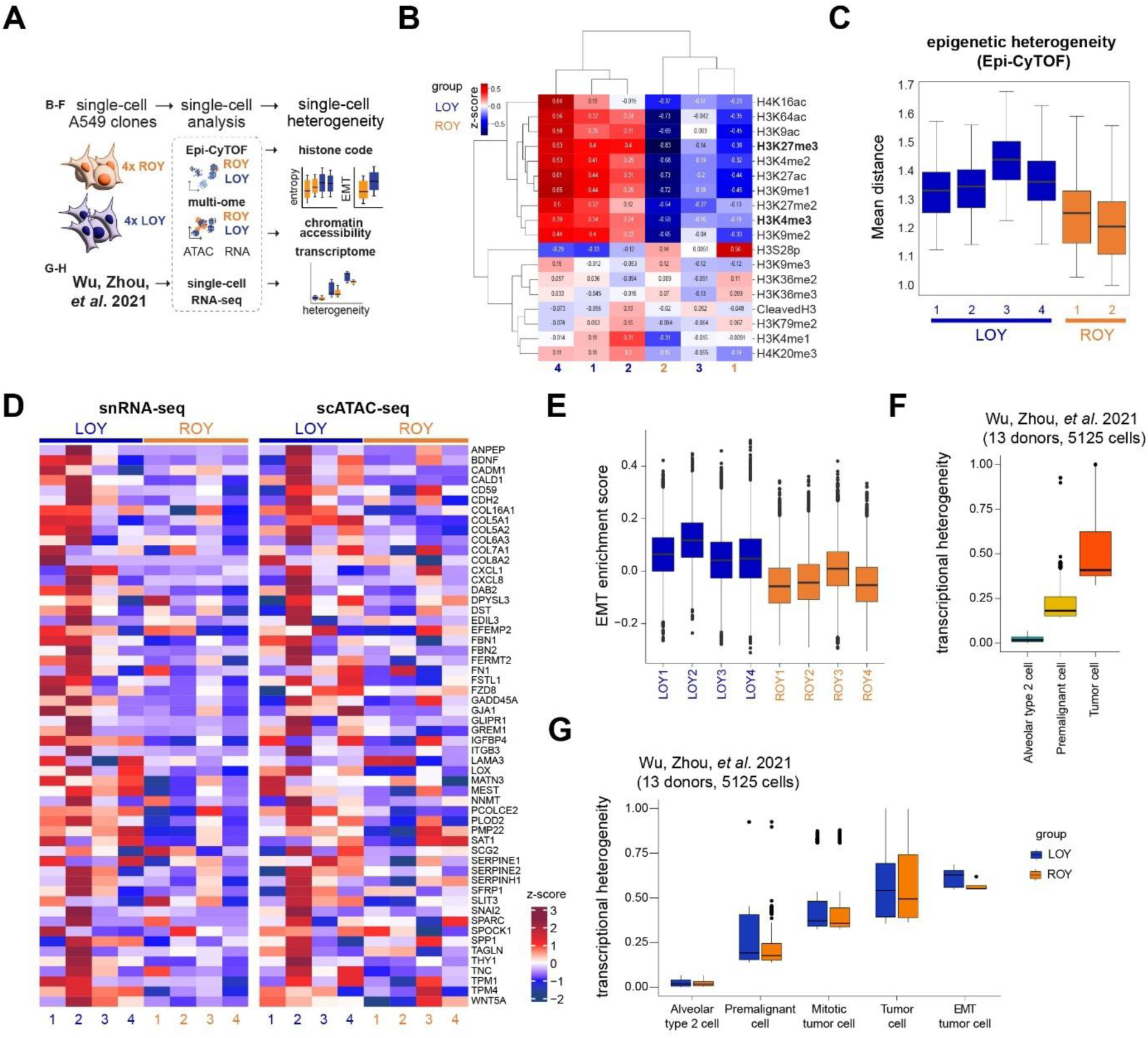
Single-cell profiling reveals LOY-driven epigenetic heterogeneity and transcriptional plasticity. **(A)** Experimental workflow for single-cell profiling of isogenic A549 ROY and LOY clones using Epi-CyTOF and scMultiome, alongside clinical data reanalysis^26^. **(B)** Heatmap of Epi-CyTOF histone modification measurements. Values are z-scored across clones; blue: lower-than-average; red: higher-than-average levels. Hierarchical clustering was applied to histone marks and samples. **(C)** Boxplot showing the epigenetic heterogeneity metric derived from multidimensional Epi-CyTOF data. **(D)** Boxplot of EMT module scores from snRNA-seq, based on lead genes from Figure 2D. **(E)** Heatmaps of lead EMT gene expression (snRNA-seq, left) and corresponding chromatin accessibility at gene promoters (scATACseq, right). **(F)** Boxplot of transcriptional heterogeneity (QuoTHiC) in premalignant and tumor cells compared to normal AT2 cells in a cohort from the LuCA dataset^26,37^. **(G)** Boxplot comparing QuoTHiC scores across cell types and ROY/LOY status from the cohort shown in (F).

At the transcriptional level, LOY clones showed loss of Y-linked gene expression without compensatory upregulation of X-linked homologs (**Figure S4C**). A549 clones were clustered separately by LOY/ROY identity in snRNA-seq, scATAC-seq, as well as in an integrated low-dimensional representation (**Figure S4D-E**). LOY clones displayed elevated yet heterogeneous EMT scores, with LOY2 showing the strongest induction of EMT genes (**Figure 4D-E**). Consistent with these transcriptional changes, scATAC-seq confirmed increased and highly variable promoter accessibility at EMT loci in LOY cells, again most pronounced in LOY2 (**Figure 4E**). Locus-level inspection confirmed enhanced accessibility at the promoters of *THY1*, *LOX*, *CDH2*, and *ANPEP* in LOY relative to ROY clones (**Figure S4G–J**), supporting a link between chromatin remodeling and activation of EMT programs.

Finally, to investigate the impact of LOY on cell state diversity in patients, we re-analyzed single-cell RNA-seq data from the LuCA^26^ dataset, specifically focusing on the cohort with the highest tumor cell recovery^26,37^ (13 LUAD donors; **Figure S4K**). We observed that transcriptional heterogeneity generally increased in premalignant and tumor cells compared to normal alveolar type 2 cells (**Figure 4F**). To further dissect this, we excluded low-quality cells based on autosome gene counts (**Figure S4L**) and classified the remaining population into ROY and LOY cells using a published random forest classifier^17^. We found that transcriptional heterogeneity was enriched in LOY cells with EMT features compared to ROY-EMT tumor cells (**Figure 4G**). Together, these results support a model where LOY induces a plastic and highly heterogeneous epigenetic state that facilitates phenotype diversification.

### LOY promotes adaptation to metabolic and genotoxic stress

We next assessed the *in vitro* functional consequences of LOY-induced plasticity and heterogeneity (**Figure 5A**). Under basal conditions, LOY and ROY clones showed comparable proliferation rates (**Figure S5A**), indicating that LOY does not confer a proliferative advantage, in line with recent observations in bladder cancer^20^. However, LOY clones showed a trend toward increased proliferation under acute metabolic stress induced by 24 hours of glucose or glutamine deprivation (**Figure S5A**). Parallel transcriptome and proteome profiling revealed that nutrient deprivation further amplified EMT programs in LOY cells. Moreover, glucose deprivation specifically enriched for oxidative phosphorylation and hypoxia signatures in LOY cells (**Figure 5B-C**, **Figure S5B-D**).

**Figure 5:**
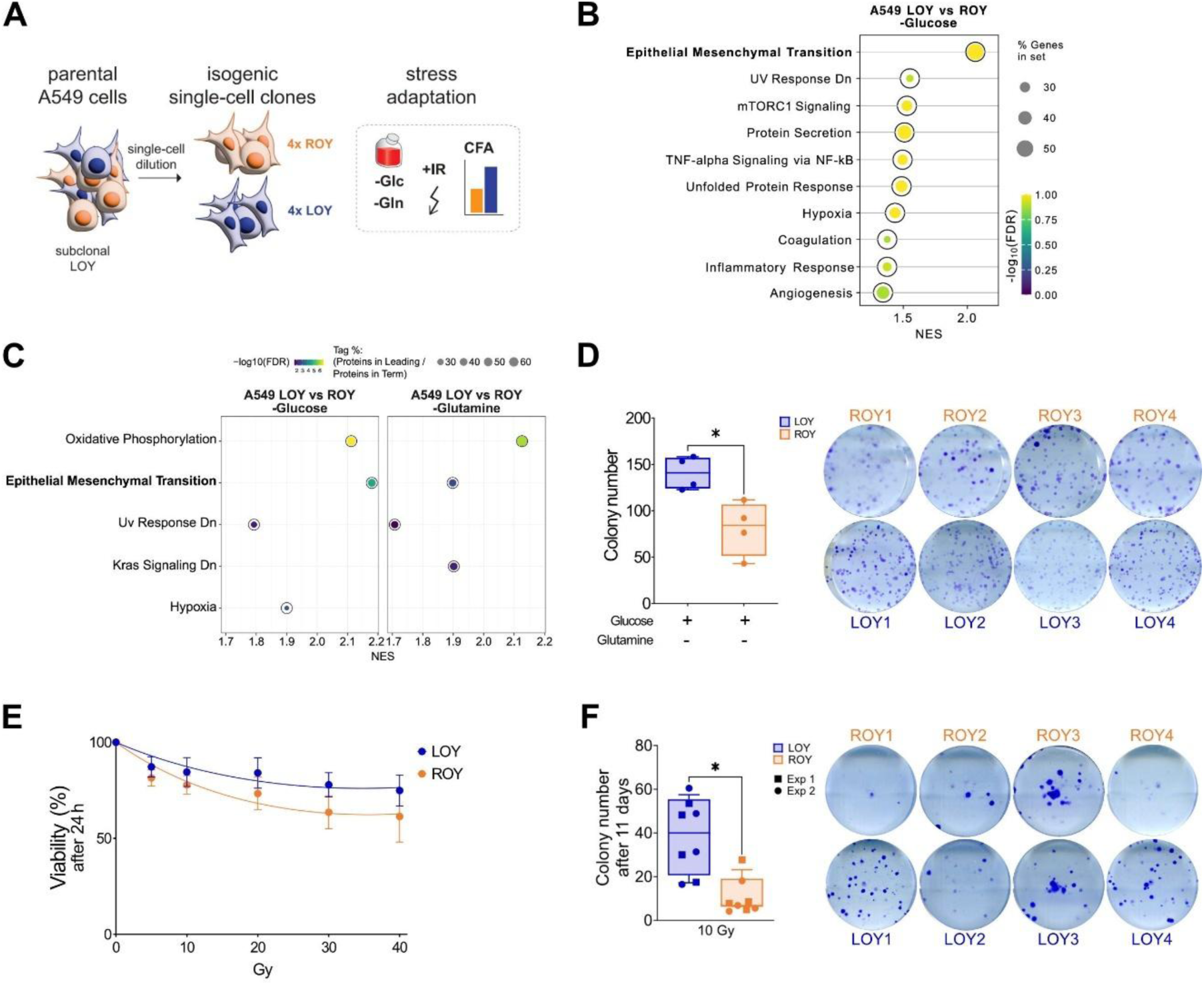
LOY promotes adaptation to metabolic and genotoxic stress. **(A)** Schematic of the *in vitro* phenotypic characterization of isogenic A549 ROY and LOY clones. **(B)** Dot plot showing the top 10 Hallmark gene sets with FDR<0.25 from preranked GSEA of RNA-seq data comparing A549 LOY and ROY under glucose deprivation. X-axis: normalized enrichment score (NES); y-axis: gene sets. Dot size indicates gene count; color represents -log_10_(FDR). (**C)** Dot plot showing significant Hallmark gene sets from preranked GSEA of global proteome data comparing LOY vs. ROY clones under glucose (left) and glutamine (right) deprivation. Dot size indicates protein count; color represents -log_10_(FDR). **(D)** Colony formation capacity under glutamine deprivation. Left: Boxplot of mean colony number (n = 3 technical replicates/clone). Right: Representative crystal violet-stained images. **(E**) Normalized dose-response curves of four LOY (blue) and four ROY (orange) clones assessed 24 hours post-irradiation (0-40 Gy). Viability was measured in triplicate using CellTiter-Blue and fitted using non-linear regression. **(F)** Clonogenic survival following 10 Gy. Left: boxplots showing mean colony counts across two independent experiments (n=3 replicates/clone/experiment). Right: representative images from experiment 1. Note: For all boxplots, the center line represents the median, box limits represent upper and lower quartiles, and whiskers represent minimum and maximum values. Statistical significance was assessed using an unpaired t-test (α=0.05).

Additionally, in clonogenic assays, LOY cells formed significantly more colonies than ROY cells, consistent with the upregulation of CD90 (**Figure 2H**, **Figure S5E**), indicating an increased stem-cell capacity. This difference was further enhanced under glutamine deprivation, a hallmark of poorly vascularized tumor microenvironments^38^, suggesting superior adaptation of LOY cells to metabolic stress (**Figure 5D**, **Figure S5E**).

LOY clones also exhibited intrinsic resistance to genotoxic stress. Following exposure to ionizing radiation (0-40 Gy), LOY cells displayed higher viability and a sustained growth advantage over five days compared to ROY cells (**Figure 5E**, **Figure S5F**). Clonogenic survival post-irradiation (10 Gy) was significantly higher in LOY clones. While colony size remained comparable, LOY cells showed a slight decrease in size (**Figure 5F**, **Figure S5G**). No differences were observed in non-irradiated controls (**Figure S5H**).

Altogether, these results demonstrate that rather than driving intrinsic proliferation, LOY confers a selective advantage by increasing the clonogenic potential and resilience to metabolic and genotoxic insults. This suggests that LOY-driven plasticity promotes tumor cell adaptability, allowing them to survive and outcompete ROY cells within hostile and nutrient-deprived microenvironments.

### LOY facilitates engraftment and metastatic dissemination

To evaluate the *in vivo* consequences of LOY in tumor cells, we analyzed two metastatic xenograft models, using A549 cells containing subclonal LOY (**Figure 6A**). First, we assessed the frequency of LOY in lung tumors formed after tail-vein injection of parental A549 cells using chromogenic *in situ* hybridization (CISH) with specific probes for the X and Y chromosomes. Across four animals, we observed that most lung tumors were predominantly composed of LOY cells (**Figure 6B**). In three out of four mice, LOY tumors represented >85% of the total tumor burden, with one animal showing 100% LOY tumor cells in the 19 tumors analyzed (**Figure 6B-C**). While we observed some intra-animal heterogeneity, including ROY and mixed LOY/ROY tumors, these results suggested a clear selective advantage for LOY cells during lung colonization.

**Figure 6:**
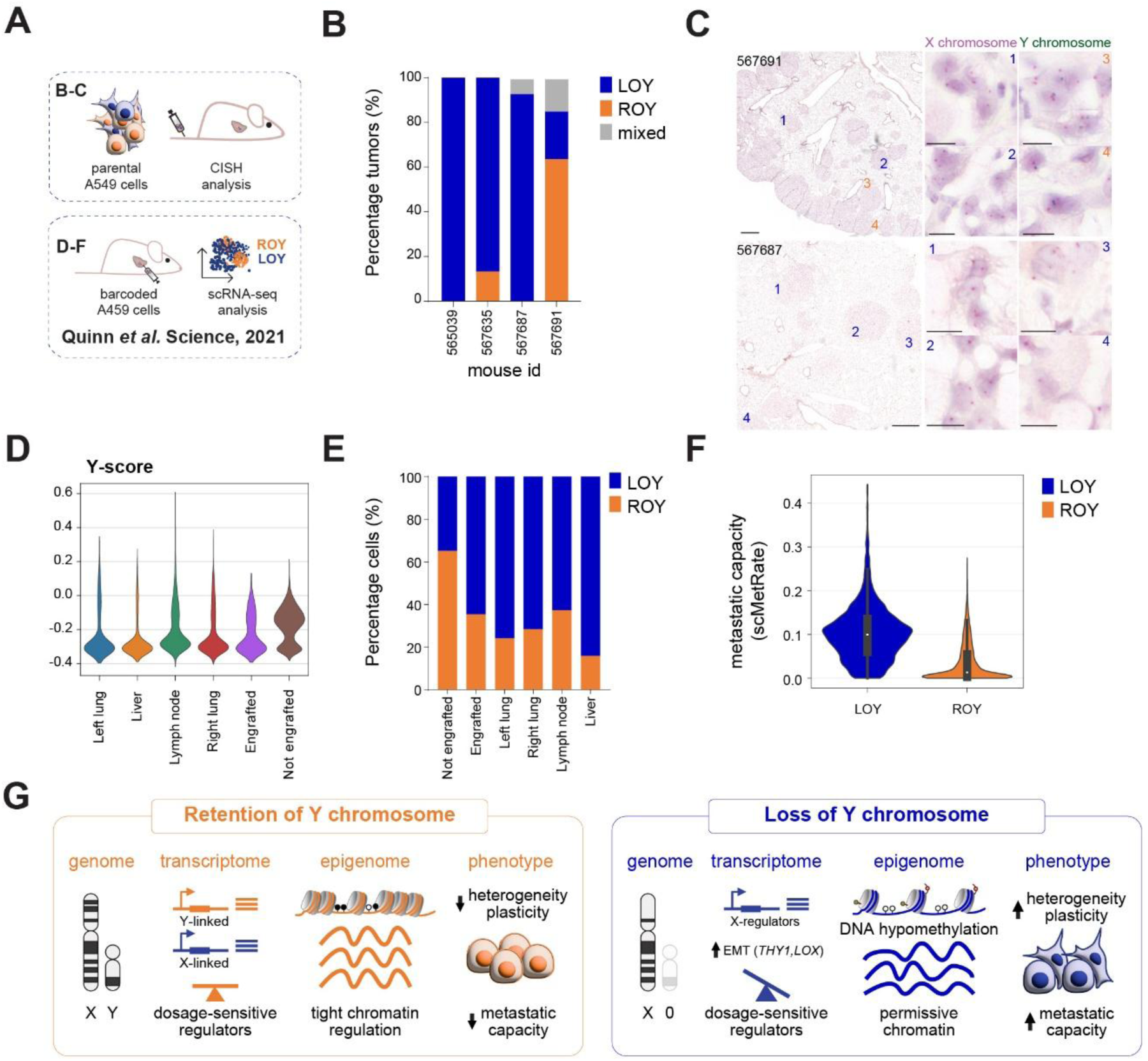
LOY facilitates engraftment and metastatic dissemination. **(A)** Experimental workflow for *in vivo* LOY characterization tail-vein (parental) and orthotopic (barcoded)^39^ xenograft models. **(B)** Quantification of LOY frequency in lung tumors via Chromosome X and Y Chromogenic *In Situ* Hybridization (CISH). Stacked bar plots represent the percentage of ROY (orange), LOY (blue), or mixed (grey; co-occurrence of LOY and ROY populations) tumors per animal. **(C)** Representative CISH visualization of LOY and ROY lung tumors. Left: low-magnification overview of tumor-bearing lungs (scale bar: 500 µm). Right: high magnification zoom into the indicated regions (scale bar: 10 µm). Probes: X chromosome (magenta), Y chromosome (green). (**D**) Violin plots showing the Y-score distribution for cells harvested from pre-implantation (not-engrafted) pools, engraftment, and metastasis sites. Data reanalyzed from Quinn *et al.*^39^. **(E)** Stacked bar plot showing the percentage of LOY (blue) and ROY (orange) cells in each anatomical compartment. **(F)** Violin plot comparing the metastatic potential of LOY vs. ROY cells based on the scMetRate metric^39^. **(G)** Working model: The Y chromosome acts as an epigenetic gatekeeper. Upon LOY, EMT gene expression and promoter hypomethylation induce plasticity and heterogeneity, promoting adaptation and metastasis.

To validate these findings in a spontaneous metastasis model and dissect the clonal dynamics driving this expansion, we re-analyzed spatially-resolved single-cell clonal tracing and scRNA-seq data from a published orthotopic xenograft model of barcoded A549 cells^39^ (**Figure 6A**). Unlike the tail-vein method, this approach allowed us to track LOY-clone dynamics throughout the entire metastatic process by comparing the abundance of LOY and ROY cells in pre-implantation cell pools, primary tumors, and distant metastases.

UMAP projection of engrafted A549 cells did not reveal clear segregation by engraftment capacity or metastatic site (**Figure S6A-C**). However, after computing Y-, X-, and housekeeping scores from the scRNA-seq data, we observed a unique, bimodal distribution of the Y-score across the entire population (**Figure 6D**), whereas X- and housekeeping scores remained uniformly distributed (**Figure S6D**). Using a Y-score threshold to classify single cells (**Figure S6E**), we observed a progressive enrichment of LOY populations during metastatic dissemination (**Figure 6E**, **Figure S6F**). While non-engrafted cells were predominantly ROY, engrafted primary tumors showed increased LOY frequency (**Figure 6E**). This enrichment was most pronounced in distant metastases, particularly in the liver (84.2% LOY). Consistent with this, metastatic potential (measured by the scMetRate metric)^39^ was increased in LOY and negatively correlated with Y-scores (**Figure 6F**, **Figure S6G**), indicating that cells lacking the Y chromosome disproportionately contribute to metastatic dissemination and outgrowth.

To further resolve LOY-clone dynamics from primary tumors to metastases, we leveraged the integration barcode (intBC)-enabled clonal-tracing labels^39^ to examine the metastatic capacity of distinct LOY- and ROY-enriched clones (defined as clones where ≥80% of cells were classified as LOY or ROY, respectively). Clones were then stratified into highly metastatic, weakly metastatic, or non-metastatic phenotypes using the categories defined in the original study^39^ (**Figure S6H**). Notably, LOY clones were significantly enriched for metastatic phenotypes, with 96% classified as highly or weakly metastatic compared to only 75% of ROY clones (**Figure 6F**, **Figure S6H**). This difference corresponds to a 21% increase in metastatic capacity for LOY cells (Fisher’s exact test, p = 0.037). Together, these data support a model in which LOY confers a selective advantage during engraftment and metastatic colonization, ultimately driving metastatic burden.

## Discussion

Loss of the Y chromosome (LOY) is among the most frequent somatic alterations in male malignancies^15–17^, yet for decades it was considered a neutral by-product of genomic instability rather than a driver of tumor cell evolution^8,16^. Recent studies have begun to challenge this view, implicating LOY in immune evasion in bladder cancer^20^, as well as in the regulation of cancer-testis antigens and remodeling of the tumor immune microenvironment in lung cancer^18^. Moreover, LOY in tumor and T cells is associated with diminished survival^17,20^, and thus, accumulating evidence across different tumor types suggests context-dependent functional consequences of LOY^8,19,22^. Still, the tumor cell-intrinsic effects of LOY have remained largely unexplored.

Our study identifies LOY as a driver of epithelial-to-mesenchymal transition (EMT) and cellular plasticity in lung adenocarcinoma (LUAD). Across primary tumors and isogenic clones, LOY was consistently associated with the activation of EMT-associated transcriptional programs. EMT is a key facilitator of lung cancer dissemination, enabling migration, immune evasion, and metastatic colonization. Specifically, LOY clones exhibited significant upregulation of EMT genes, including *THY1* (CD90), a well-established marker of cancer stem cells and poor prognosis in lung cancer^40–42^, and *LOX*, an enzyme involved in extracellular matrix remodeling and metastatic progression^43,44^.

Our findings establish EMT as a central biological consequence of Y chromosome loss and provide a mechanistic framework linking LOY to metastasis and poor clinical outcome in male lung adenocarcinoma. Moreover, these findings were confirmed in an independent study by Dorsch *et al.* in pancreatic adenocarcinoma (submitted in parallel), further supporting the role of LOY as a driver of EMT and tumor cell plasticity.

Importantly, EMT activation was heterogeneous, spanning partial and intermediate EMT states rather than a uniform epithelial-to-mesenchymal switch, expanding the diversity of intermediate and mesenchymal-like states from which aggressive phenotypes are preferentially selected during tumor progression and metastasis. This observation aligns with emerging models of EMT as a continuum of metastasis-competent phenotypes rather than a binary process, and with recent models in which metastasis emerges from phenotypically diverse subpopulations rather than from a deterministically reprogrammed single-state^45,46^.

Functionally, rather than accelerating proliferation under baseline conditions, LOY increases cellular plasticity and the capacity to adapt to metabolic and genotoxic stress. Because of their enhanced plasticity, we found that LOY cells are more resistant to ionizing radiation. Moreover, *in vivo*, this plasticity provides a selective advantage during tumor engraftment and metastatic dissemination.

A central insight of our study is the identification of the Y chromosome as an epigenetic “gatekeeper” that preserves cell identities and maintains homeostasis across epigenetic, transcriptional, translational, and proteomic networks^2,3^ (**Figure 6G**). We propose that LOY leads to loss of Y-linked dosage-sensitive regulators, which in turn destabilizes the chromatin landscape. A hallmark of this destabilization is the widespread DNA hypomethylation at regulatory elements, an observation also confirmed in patients^24^, together with altered patterns of histone posttranslational modifications. DNA hypomethylation and H3K4me3 levels were enriched at the promoters of EMT-associated genes, including *THY1* and *LOX*, lowering the epigenetic barrier for the activation of aggressive transcriptional programs.

However, beyond specific loci, the most striking feature of LOY cells was an increase in cell-to-cell variability rather than uniform global changes. Our single-cell analyses revealed that LOY clones displayed increased epigenetic heterogeneity and transcriptional plasticity compared to isogenic ROY clones. In our model, the Y chromosome acts as a stabilizer of cellular states, and its loss increases stochastic fluctuations in epigenetic and transcriptional states (**Figure 6G**). In this framework, cell state switching does not primarily arise through active reprogramming, but rather through the Darwinian selection of cells with higher fitness. As these cells are present within a pool of heterogeneous cell states, the tumor can mount a rapid response to selective pressures.

Thus, LOY acts as a lineage-plasticity amplifier, driving malignancy through epigenetic diversification rather than classical oncogenic signaling, providing a mechanistic framework linking LOY to poor outcome in male LUAD. Future work will be required to define the specific Y-linked regulators and signaling circuits responsible for these effects, and to determine whether LOY-associated heterogeneity creates therapeutic vulnerabilities or predicts response to standard-of-care treatment.

### Limitations of the study

We acknowledge that our in vitro experiments relied on a single lung cancer cell line. Although we analyzed four independent sublines per condition to rule out clonal artifacts, the cell-line-specific genetic context of LOY may still influence the phenotype. We validated our key findings using primary bulk and scRNA-seq data from LUAD patients and independent xenograft models^39^. Moreover, parallel findings in pancreatic ductal adenocarcinoma (PDAC) models and primary samples (Dorsch et al., submitted) strongly suggest that the observed EMT phenotype is a conserved mechanism across different tumor entities rather than an A549-specific artifact.

Additionally, we did not identify the specific Y-linked gene(s) whose haploinsufficiency drives this cellular plasticity. Future work is required to pinpoint these drivers, though *KDM5D* represents a compelling candidate. As a H3K4me3 histone demethylase^3^, KDM5D might influence DNA methylation similarly to its X-linked homolog KDM5C^47^. Its loss partially phenocopies LOY effects in bladder cancer and acute myeloid leukemia mouse models^20,48^. In addition, recent work by Chen *et al.*^49^ indicates that *KDM5D* knockout (KO) in A549 cells promotes tumorigenesis and lung metastasis via p38α signaling. Nevertheless, the potential contribution of other Y-linked genes will require further gain- and loss-of-function studies.

## Methods

### Study approval

The protocol for tissue collection was approved by the ethics committee of the University of Heidelberg (biobank vote: S-270/2001; and study protocol: S-154/2018). The experiments were conducted in accordance with the principles outlined in the WMA Declaration of Helsinki and the Department of Health and Human Services’ Belmont Report. All patients provided written informed consent before being included in the study and remained anonymous throughout the study.

### Patient samples

Fresh tumor and paired distal lung samples from lobe resection surgeries at Thoraxklinik-Heidelberg were provided by Lung Biobank Heidelberg, a member of the accredited NCT Tissue Bank, the BioMaterialBank Heidelberg (BMBH), and the platform biobanking of the German Center for Lung Research (DZL). All patients provided written informed consent for the use of their specimens for research purposes (see study approval). Strict patient inclusion criteria were established at the beginning of the prospective tissue collection. To avoid the effects of acute inflammation caused by smoking, all donors stopped smoking at least three months before donating. None of the donors had received chemotherapy or radiation therapy within four years before surgery. Additionally, we collected lung function results based on forced expiratory volume in one second (FEV_1_) and the FEV_1_/forced vital capacity (FVC) ratio, as well as medical history for each patient to best characterize the included samples (**Table 1**). All histopathological diagnoses were made according to the current WHO classification for lung cancer by at least two experienced pathologists. The tumor stage was designated according to the 8th edition of the UICC tumor, node, and metastasis. In total, paired distal lung and lung adenocarcinoma tissue from four former-smoker donors were used for whole-genome sequencing (flash-frozen material, see below) and single-cell RNA-seq (cryopreserved samples).

### Whole-genome sequencing of tissue samples

DNA from the tumor and paired distal lung samples was provided by the Lung Biobank Heidelberg. Tissues were snap-frozen within 30 minutes after resection and stored at -80°C until further processing. For DNA isolation, 10 - 15 tumor cryosections (10 - 15 μm each) were prepared for each patient. Frozen cryosections were homogenized with the TissueLyser mixer-mill disruptor (2 x 2 min, 25 Hz, Qiagen, Hilden, Germany). DNA was isolated using the AllPrep DNA/RNA Mini Kit (Qiagen) according to the manufacturer’s instructions. DNA concentration was measured using the Qubit™ dsDNA BR Assaykit (Invitrogen). Whole-genome sequencing libraries were generated using the TruSeq Nano DNA Kit (Illumina) at the DKFZ NGS core facility. Libraries were sequenced on a NovaSeq 6000 system using the S4 v1.5 flow cell with paired-end 150 bp reads, and a coverage of 80x for each tumor sample, and 40X for the distal lung samples.

### Whole-genome sequencing - analysis

WGS data were aligned to the hg38 reference genome using the Burrows-Wheeler Aligner^50^. The WGS data processing downstream of the aligned BAM files was performed using a Nextflow pipeline: https://github.com/CompEpigen/wf_WGS. As part of this pipeline, the Manta workflow was run to detect structural variants (SV)^51^. Control-FREEC was used to call copy number variations (CNV)^52^. For donors, we performed an additional step for calling SNVs and indels using Mutect2 and matched normal samples and a list of 75 driver genes for lung adenocarcinoma from intogen.org (**Table 4**). Driver mutations, CNVs, and SVs are reported in **Tables 2, 5, and 6**. WGS data were used to demultiplex the scRNA-seq data from the four male donors. Plots were generated using the software Figeno^53^.

### TCGA somatic mutation data

Somatic mutation calls were obtained from the TCGA-PanCanAtlas Publications MC3 mutation dataset (mc3.v0.2.8.PUBLIC.maf.gz). To ensure a single representative tumor sample per patient, TCGA barcodes were truncated to the first 12 characters to define patient-level ID, and duplicates were removed. Only male primary tumor samples were included, and genes with mutation recurrence in ≥ 5 samples were retained. LOY status was assigned following Qi *et al.*^15^ using TCGA WGS/WES-based Y chromosome copy number states^15^. Samples with gain plus loss were excluded, while LOY-related states were grouped as LOY and wild-type or gain states as ROY. TCGA samples were jointly analyzed with the four male in-house samples generated in this study. Comparative analyses between LOY and ROY groups were conducted using maftools (v2.26.0). Mutation frequencies of previously reported lung adenocarcinoma driver genes were compared through the coOncoplot function, based on a curated list of 75 driver genes from the IntOGen cancer driver database (downloaded on Dec. 4th, 2023; six LUAD cohorts comprising over 1,200 tumors). Enrichment between LOY and ROY groups was assessed using Fisher’s exact test (mafCompare) and visualized as forest plots (forestPlot).

### Cryopreservation of lung parenchyma and tumor samples

Freshly excised pieces of paired distal lung tissue and lung adenocarcinoma tissue were transported in a CO₂-independent medium (Thermo Fisher Scientific) supplemented with 1% bovine serum albumin (BSA) (Carl Roth), 1% penicillin/streptomycin (Fisher Scientific), and 1% amphotericin B (Sigma). The tissue was processed as previously published^54–57^. Briefly, the tissue pieces were inflated with cold HBSS (Fisher Scientific) supplemented with 1% BSA (Carl Roth), 2 mM EDTA (Fisher Scientific), 1% amphotericin B, and 1% penicillin/streptomycin (Fisher Scientific). Exemplary pieces of distal normal lung tissue and tumors were collected for histological analysis (see above). The pleura was carefully removed from the distal lung tissue. Each tissue compartment (distal lung, pleura, and adenocarcinoma) was minced into 4 × 4 mm pieces using a sterile razor blade and scissors. 15-20 pieces were cryopreserved separately in cryotubes (Sarstedt) with 1.4 mL of ice-cold freezing medium consisting of 70% high-glucose DMEM GlutaMAX™ (Thermo Fisher Scientific), 10% DMSO (Carl Roth), and 20% FBS (GE Healthcare). The tubes were flipped to distribute the medium within the tissue pieces and kept on ice for fifteen minutes. The tubes were placed into Mr. Frosty containers (Thermo Fisher Scientific) and transferred to –80°C to ensure a cooling rate of 1°C/min. For long-term storage, the samples were kept in liquid nitrogen.

### Tissue dissociation, sorting, and sample pooling for scRNA-seq

To minimize technical bias, all the samples were processed in parallel. Paired cryopreserved distal normal lung and tumor tissues from four male donors were thawed for two minutes in a water bath at 37°C. The thawed tissue samples were collected in 50 mL Falcon tubes (Neolab Migge) and washed with HBSS (Fisher Scientific) supplemented with 1% BSA (Carl Roth), 2 mM EDTA (Fisher Scientific), 1% amphotericin B, and 1% penicillin/streptomycin (Fisher Scientific). The samples were minced into smaller pieces before undergoing mechanical and enzymatic dissociation to generate single-cell suspensions using the human tumor dissociation kit (Miltenyi Biotec)^54,57^. Briefly, 0.5–1 g of minced tissue was added to a MACS C tube containing the enzyme mix from the human tissue dissociation kit (Miltenyi Biotec), consisting of 200 μL of enzyme H, 100 μL of enzyme R, and 50 μL of enzyme A in 4.5 mL of CO₂-independent medium (Thermo Fisher Scientific) supplemented with 1% bovine serum albumin (BSA) (Carl Roth), 1% penicillin/streptomycin (Fisher Scientific), and 1% amphotericin B (Sigma). 10 μM of the ROCK inhibitor (Y-27632) and 100 μL of DNase I were added to each tube. The tubes were tightly closed and introduced into the MACS dissociator and mechanically disrupted using program h_tumor_01, followed by a 15-minute incubation at 37°C on a rotator; h_tumor_01, plus 15 minutes at 37°C on a rotator; h_tumor_02, and 15 minutes at 37°C on a rotator for a final enzymatic dissociation and a last mechanical shearing using the program h_tumor_02. The samples were pipetted up and down to help disaggregate them. Finally, the enzymatic reaction was stopped by adding 20% FBS (Gibco, Thermo Fisher Scientific), and single cells were collected by sequential filtering through 100 μm, 70 μm, and 40 μm cell strainers (BD Falcon). Single-cell suspensions were centrifuged for 8 minutes at 4°C and 200g, resuspended in 4 mL ACK lysis buffer (Millipore Sigma), and incubated for 3 minutes at room temperature to lyse erythrocytes. The reaction was stopped by adding 35 mL of HBSS/1% BSA/2 mM EDTA/1% amphoB/1% pen/strep. Cells were centrifuged for 8 minutes at 4°C and 200g, resuspended in 0.5 mL HBSS/1% BSA/2 mM EDTA/1% amphoB/1% pen/strep, and counted. Fc receptors were blocked with human TruStain FcX (BioLegend) for 30 minutes on ice. Immune and epithelial cells were labeled with CD45-APC Cy7 (BioLegend) and EPCAM-PE (CD326, Affymetrix eBioscience) antibodies for 30 minutes in the dark at 4°C following the manufacturer’s instructions. Stained samples were washed, resuspended in 0.5 mL HBSS/1% BSA/2mM EDTA/1% amphoB/1% pen/strep, and added to FACS tubes with 40 μm cell strainer caps. To discriminate between live and dead cells, SyTOX blue was used as recommended (Thermo Fisher Scientific). We sorted live, single-cell-gated CD45^+^ and CD45^neg^ cells using a FACSAria II cell sorter (BD Biosciences). The sorted CD45^+^ and CD45^neg^ cells derived from paired distal normal lung and lung adenocarcinoma samples were used for single-cell RNA sequencing analysis (10X Genomics, Single Cell 3′ Gene Expression v3.1). Approximately 5,000 cells per donor, derived from the same tissue type (distal normal or tumor) and sorted compartment (CD45^+^ and CD45^neg^), were combined (20,000 cells in total from 4 donors per well) and loaded in each well of the Chromium Next GEM Chip G (10X Genomics). FlowJo software (Tree Star) and Kaluza (Beckman Coulter) were used to analyze and plot the FACS results.

### Single-cell RNA-seq library preparation

5,000 CD45^+^ or CD45^neg^ sorted cells from cryopreserved distal normal lung or lung adenocarcinoma of four male donors were pooled and loaded per well of the Chromium Next GEM Chip G (10X Genomics) and loaded into the 10X Genomics Chromium controller according to the manufacturer’s protocol. Reverse transcription, cDNA amplification, and the subsequent library preparation from 250 ng of cDNA were performed following the 10X Genomics Single Cell 3′ gene expression v3 protocol. 10nM of the finished libraries were pooled and sequenced using Illumina NovaSeq 6000 instrument with an S4 flow cell, generating approximately 1,450 million 100bp paired-end reads.

### Single-cell RNA-seq analysis

As described above, the scRNA-seq samples were generated separately for CD45^+^ and CD45^neg^ fractions from distal normal and tumor regions, yielding four sample types (CD45^+^ distal normal, CD45^neg^ distal normal, CD45^+^ tumor, and CD45^neg^ tumor), with each sample pooled from four male patients. For each sample, preprocessing was performed using CellRanger (v6.1.1) followed by scCancer (v2.3.0), which includes cell- and gene-level quality control. Quality control cutoffs were automatically determined by scCancer based on total UMI counts, the number of detected genes, and the proportion of mitochondrial transcripts. For each pooled sample, donor demultiplexing was performed using Souporcell (v2.4.0) to identify variants per cell, followed by cell-level allele counting and genotype-based cell clustering with k = 4 clusters. Doublets were identified using troublet (v2.4; a tool in Souporcell pipeline) and excluded. Inferred cluster genotypes were matched to germline SNP calls from whole-genome sequencing data using Picard (v2.25.1) GenotypeConcordance function to assign donor identities. Batch effects were mitigated using the NormalMNN module in scCancer across samples. Cell state scores, including infercnv-based malignancy, stemness, and cell cycle scores, were computed using scCancer, while Y score and EMT score were calculated using Seurat’s function AddModuleScore (**Figure S1G and S1L; Table 7**).

### Cell type annotation and tumor cell identification

Cells were initially clustered using the Leiden algorithm in latent space. Epithelial compartments were manually annotated based on canonical marker genes, while automated reference-based annotation for all cells was performed with Azimuth snakemake workflow (v0.3.1) using Human Lung Cell Atlas (HLCA) as the reference. Cells with inconsistent annotations between manual epithelial labels and Azimuth predictions were excluded from further analysis. Tumor versus non-tumor epithelial status was inferred using the malignancy scoring module in scCancer, which incorporates copy number variation (CNV) profiles generated by inferCNV.

### Survival analysis: Kaplan-Meier Plot

Kaplan-Meier survival analysis was performed using the kmplot.com online tool^58^. The analysis was restricted to male LUAD patients from the mRNA gene chip dataset. The mean expression of five Y-linked genes (*RPS4Y1*, *DDX3Y*, *UTY*, *KDM5D*, *EIF1AY*) per patient was used to stratify patients into low- and high-expression groups using the tool’s auto-select best cutoff. For a second approach, the mean expression of 13 Y-linked genes^29^ (*RPS4Y1*, *ZFY*, *PCGH11Y*, *TBL1Y*, *PRKY*, *USP9Y*, *DDX3Y*, *UTY*, *TMSB4Y*, *NLGN4Y*, *TXLNGY*, *KDM5D*, *EIF1AY*) per patient was used to stratify patients into low- and high-expression groups using patients in quartile 1 and quartile 4. Biased arrays were excluded from the analysis. Survival differences between groups were evaluated using univariate Cox proportional hazards regression.

### TCGA bulk RNA-seq GSEA

Bulk RNA-seq count matrix was obtained from TCGA via TCGAbiolinks (v2.24.3) using “STAR - Counts” workflow and filtered for male LUAD patients and full LOY status (n=75) or full ROY status (n=110) based on^15^. Fractional or partial LOY was excluded. The expression matrix was filtered to include genes with at least 10 counts in at least 25 samples within the LOY or ROY. Differential expression analyses between LOY and ROY LUAD patients were performed using pyDESeq2 (v0.4.10; Python v3.11.9)^59^. Statistical significance was assessed using Wald test and p-values were adjusted for multiple testing using the Benjamini-Hochberg procedure. Gene set enrichment analysis was performed using the prerank implementation in GSEApy (v1.1.3; Python v3.12.5)^60^. Genes were ranked by log2FC from differential expression analysis. Enrichment was assessed using the MSigDB Hallmark gene set collection (v7.1). Enrichment significance was determined using 1,000 gene set permutations, and enrichment results were corrected for multiple testing using the false discovery rate.

### Single-cell lung cancer atlas (LuCA)

The LuCA^26^ core data set was downloaded from cellxgene website. Tumor cells were extracted from primary tumor regions of lung adenocarcinoma of male patients. Quality control of cells was performed using a cross-point threshold defined by the Y gene and autosomal gene detection rates to exclude false predictions of LOY cells caused by high dropout. Cells with zero detected Y-linked genes, but sufficiently high autosomal gene detection rates above the defined threshold, were retained for downstream analysis. Next, LOY status at the single-cell level was computationally predicted using a published Random Forest model^17^. Tumor cells from female lung adenocarcinoma patients were used as a positive control for LOY. The expression of Y-linked genes in three groups - male LOY, male ROY, and female - was visualized by Seurat (v5.2.1) dotplot. Gene set enrichment analysis was performed on male tumor cells using the GSEApy^60^ package (v1.1.11), with the Human Molecular Signatures Database (MSigDB_Hallmark_2020) and the “signal_to_noise” method with 1,000 permutations. The top 8 pathways (NES>0) that are enriched in the LOY cells were displayed. Cell downsampling was applied before the computation of cell state scores (malignancy, Y, stemness, cell cycle, and EMT scores) to control for differences in cell numbers between groups, retaining a maximum of 1,500 cells per cell type. Transcriptomic heterogeneity was assessed within the Wu-Zhou cohort^26,37^, which represents the major study enriched for tumor cells with EMT features. Heterogeneity was quantified using silhouette scores as implemented in the QuoTHiC package (github.com/CompEpigen/quothic). The top 2,000 highly variable genes were selected, and the first 50 principal components were used for UMAP construction, followed by distance calculations. For each cell *i*, the silhouette score is defined as *s*(*i*)=(*b*(*i*)−*a*(*i*))/max{*a*(*i*),*b*(*i*)}, where *a*(*i*) is the average distance to cells within the same group and *b*(*i*) is the minimum average distance to cells in other groups. To ensure that higher values indicate greater heterogeneity, the score was multiplied by −1, and the resulting values were min–max scaled to the range [0,1].

### Cell culture assays

#### General cell culture

A549 cells were purchased from Cytion GmbH. They were authenticated using SNP typing and regularly tested for contamination using the services from Multiplexion GmbH (Heidelberg, Germany). The cells were cultured in RPMI 1640 Medium with L-Glutamine (Gibco) supplemented with 10 % FBS Superior (Sigma-Aldrich) and 100 units/ml Penicillin and 100 µg/ml Streptomycin (Sigma-Aldrich) in 10cm cell culture dishes (Greiner Bio-One). Cells were maintained in culture for one week after thawing before being used for experiments. Cells were subcultured before reaching 80% confluency. Cultures were not maintained for longer than two months. Fresh vials were thawed to initiate new culture as needed. Medium was always refreshed one day before harvesting cells for experiments. For subculturing and harvesting, cells were incubated with TrypLE Express (Gibco) for 5 minutes to detach and afterwards centrifuged at 500xg for 5 minutes. Cell count and viability were assessed using the Solution 13 AO•DAPI Staining Reagent (ChemoMetec) and the NucleoCounter® NC-250™ (ChemoMetec) according to the manufacturer’s instructions.

#### Single-cell dilution to generate LOY and ROY single-cell clones

To achieve single-cell seeding, a dilution series was performed to adjust the cell suspension to a final concentration that would theoretically result in 0.5 cells per well. The parental A549 cell suspension was seeded into a 96-well plate (TPP) accordingly. At this concentration, statistically, approximately half of the wells would receive a single cell, while the others would remain empty or contain more than one cell. After 5 days, it was recorded which wells had no colonies, one colony, or multiple colonies. After 10 days, the cells that arose from a single colony were expanded in a 24-well plate, and a fraction was used for DNA isolation. DNA isolation was performed for each clone using the Extracta DNA Prep for PCR kit (Quantabio) according to the manufacturer’s instructions. Presence or absence of the Y chromosome was confirmed by end-point PCR using a set of primer pairs targeting several Y chromosomal regions and control regions on an autosome (Key resource table). After expansion, clones were cryopreserved and stored in liquid nitrogen for long-term storage. The complete loss of the Y chromosome was confirmed using WGS.

#### ELISA-based measurement of proliferation

Cells were seeded at a density of 1×10^4^ cells per well in 96-well plates and allowed to adhere for 24 hours. Following, cells were deprived of serum and nutrients for 24 hours to synchronize metabolic states and minimize baseline proliferation. After starvation, cells were treated with three culture media conditions for 24 hours. The control group was maintained in complete RPMI-1640 medium (with L-glutamine (300 mg/L) and D-glucose (2000 mg/L)) supplemented with 10% fetal bovine serum (FBS) and 1% penicillin-streptomycin (10,000 U/mL). Experimental groups were cultured in media lacking either D-glucose or L-glutamine to assess the metabolic dependence of proliferation under nutrient-deprived conditions. Cell proliferation was quantified using the bromodeoxyuridine (BrdU) cell proliferation assay (Roche), according to the manufacturer’s instructions.

#### Colony Formation

To assess colony-forming ability under glutamine-deprived conditions, 300 cells per well were seeded in triplicate into 6-well plates in either RPMI 1640 Medium without L-Glutamine (Gibco) or RPMI 1640 Medium with L-Glutamine (Gibco), both supplemented with 10 % FBS (Sigma-Aldrich). After 5 days of incubation, the medium was replaced with fresh medium. After 10 days, cells were washed with PBS and fixed with 70% ethanol for 10 minutes at room temperature. Following fixation, ethanol was aspirated, and plates were left at room temperature for 40 minutes to allow complete evaporation of residual ethanol. Colonies were stained with 1% crystal violet solution for 10 minutes. Excess stain was removed, and wells were rinsed with tap water until the background was transparent. Colonies were manually counted using an electronic colony counter.

#### Irradiation

For the dose-response curve, cells were seeded at a density of 2,000 cells per well in a 96-well plate, in triplicate for each treatment condition and read-out. To allow cell attachment, the cells were irradiated with ionizing radiation after 24 hours at doses of 0, 5, 10, 20, 30, and 40 Gy using the MultiRad225 (Precision X-Ray). Cell viability was assessed right before irradiation and subsequently assessed every 24 hours post-irradiation using the CellTiter-Blue® Cell Viability Assay (Promega). 20 µL CellTiter-Blue reagent was added to 100 µL complete medium per well, and the mixture was incubated in the dark for 2 hours at 37 °C. Fluorescence was measured using a SpectraMax iD3 microplate reader (Molecular Devices) according to the manufacturer’s instructions. Cell viability was normalized to the untreated controls.

For the colony formation assay, 8.5×10^5^ cells were seeded into a 10 cm dish. The cells were irradiated with 10 Gy after 20 hours. Subsequently, cells were cultured for 48 hours. Afterwards, 2,000 cells were seeded into 6-well plates in triplicate from both irradiated and non-irradiated control dishes. In a second independent experiment, 200 cells were seeded into 6-well plates in triplicate for non-irradiated controls, and 2,000 cells for irradiated as before. The medium was changed every 3 days. For both experiments, after 11 days, cells were fixed with 10% formalin (Sigma-Aldrich), washed, and then stained with crystal violet and counted with an electronic colony counter as described above. Additionally, % Coverage was calculated using Fiji^61^ (ImageJ version 1.49k, National Institutes of Health, Bethesda, MD, USA), a 400 x 400 pixel circle was placed into the center of the well, the plate scan was converted to 8-bit, and the threshold was manually changed to reflect the stained area for each well. From this % area was calculated ((area covered by stain/total area of circle) × 100).

### Whole genome sequencing of A549 cells

Genomic DNA was isolated from cultured cell lines using the DNeasy Blood & Tissue Kit (Qiagen) according to the manufacturer’s instructions. Cells were harvested, and approximately 1 × 10^6^ cells were pelleted as described in the section on cell culture. DNA was eluted in 100 µL of nuclease-free water. The concentration was measured using the Qubit™ dsDNA BR Assaykit (Invitrogen). Whole-genome sequencing libraries were generated using the TruSeq Nano DNA Kit (Illumina). Libraries were sequenced on a NovaSeq 6000 system using the S4 v1.5 flow cell with paired-end 150 bp reads and 30x coverage per sample.

The alignment, processing, and plotting were performed as described for the tissue samples. The WGS Nextflow pipeline parameters were adapted to run with the WGS from A549 parental cells as a control and only use Control-FREEC and Manta. CNVs and SVs are reported in **Table 8-9**.

### Bulk RNA sequencing of A549 cells

For the bulk transcriptome analysis, 700,000 cells were seeded in standard supplemented RPMI medium (Gibco) in a 10 cm dish (Greiner) for standard growth and nutrient-deprived conditions. After 24 hours, the medium was changed to medium lacking glutamine or glucose or to fresh standard medium. After an additional 24 hours, the medium was aspirated, cells were washed twice with PBS on ice and harvested using a cell scraper. Cell pellets (approximately 5×10^6^ cells) were snap-frozen and stored at -80°C before isolation. Total RNA was isolated from cultured cell lines using the RNeasy Plus Mini Kit (Qiagen) according to the manufacturer’s instructions. The concentration was measured using the Qubit™ RNA BR Assaykit (Invitrogen). RNA integrity was determined using RNA ScreenTape (Agilent). Sequencing libraries were generated using the TruSeq Stranded mRNA Kit (Illumina). Libraries were sequenced on a NovaSeq 6000 system using the S4 v1.5 flow cell with paired-end 100 bp reads and 60-90 M reads per sample. The nf-core/rnaseq (10.5281/zenodo.1400710; version 3.14.0) was used for downstream processing, alignment to hg38 and quantification as well as quality control steps. The expression matrix was filtered to include genes with at least 10 counts in at least 3 samples within the LOY (n=4) or ROY (n=4) group. Gene-level differential expression analyses between LOY and ROY clones were performed using pyDESeq2 with default settings (v0.4.10; Python v3.11.9^59^). Statistical significance was assessed using Wald test and p-values were adjusted for multiple testing using the Benjamini-Hochberg procedure. Gene set enrichment analysis was performed using the prerank implementation in GSEApy (v1.1.3; Python v3.12.5^60^). Genes were ranked by log2FC from differential expression analysis. Enrichment was assessed using the MSigDB Hallmark gene set collection (v7.1). Enrichment significance was determined using 1,000 gene set permutations, and enrichment results were corrected for multiple testing using the false discovery rate.

### Whole-proteome profiling

#### Cell lysis and sample preparation

For basal and nutrient-deprived conditions (70% confluency), 700,000 cells were seeded in 10 cm dishes (Greiner). For the 100% confluence condition, 1×10^6^ cells were seeded. After 24 h, the medium was replaced with either standard medium or medium lacking glucose or glutamine. Following an additional 24 h, cells were washed twice with ice-cold PBS and harvested on ice using a cell scraper. Pellets were snap-frozen and stored at -80°C. Samples were thawed on ice and lysed in a buffer containing 25 mM Tris pH 7.6, 150 mM NaCl, 0.1% SDS, 1% NP40, 1% Deoxycholic acid NA-salt, 1 mM Na3VO4, 10 mM NaF, 1 µg/mL Aprotinin, 0.1 mg/mL 4-(2-Aminoethyl) benzene sulfonyl fluoride-hydrochloride, 250 U/mL Benzonase, 10 U/mL DNase and PhosSTOP (Roche: 1×Tablet/10 ml buffer). Cell lysates were incubated and rotated for 30 min at 4°C, centrifuged at 14000 rpm for 15 min at 4°C, and supernatants transferred to protein low-bind tubes. Protein concentration was determined by BCA Assay (Pierce). 5 µg of total protein per sample was used for downstream processing.

#### Protein digestion and clean-up

Automated paramagnetic bead-based single-pot, solid-phase-enhanced sample-preparation (Auto-SP3) was performed on a Bravo liquid handling platform (Agilent) as previously described^62^. Briefly, 5 µg of protein per sample was reduced with 10 mM TCEP and alkylated with 40 mM CAA (5 min at 95°C). Sera-Mag Speed Beads (A and B; Cytiva) were added to the samples, and protein digestion was performed overnight at 37°C using trypsin (1:25 enzyme:protein ratio) in 100 mM TEAB. Recovered peptides were dried via vacuum centrifugation and stored at -80°C.

#### Mass spectrometry acquisition

All samples were analyzed using an Ultimate 3000 HPLC (Thermo Fisher Scientific) coupled to an Orbitrap Eclipse Tribrid mass spectrometer (Thermo Fisher Scientific). Tryptic peptides were resuspended in 15 µl loading buffer (0.1% formic acid (FA), 2% ACN in MS-compatible H_2_O), and 2 µL were injected per sample. Peptides were loaded onto a pre-column (PEPMAP 100 C18 5 µm 0.3 mm × 5 mm, Thermo Scientific) using a loading pump at a higher flow rate. After 4 min, a valve was switched, and peptides were delivered to an analytical column (75 µm × 15 cm, packed in-house with Reprosil-Pur 120 C18-AQ, 1.9 µm resin, Dr. Maisch) at a flow rate of 5 µL/min in 98% buffer A (0.1% FA in MS-compatible H_2_O). After loading, peptides were separated using a 60 min gradient from 8% to 38% of buffer B (0.1% FA, 80% ACN in MS-compatible H2O) at a 300 nL/min flow rate. The Orbitrap Eclipse Tribrid mass spectrometer was operated in data-independent mode (DIA), with an m/z range of 350-1400. Full scan spectra were acquired in the Orbitrap at 120 000 resolution after accumulation to the set target value of 300% (100% = 1e^6^) and maximum injection time of 45 ms. DIA scans followed the full scans. 21 isolation windows were defined, with an m/z range of 411-980. Spectra were generated in the orbitrap (isolation window 1 m/z) after fragmentation using higher energy collisional dissociation (HCD) at normalized collision energy (N)CE of 28 % and acquired at 30 000 resolution after accumulation to the set target value of 1000% (100% =1e^5^) and maximum injection time of 54 ms.

#### Data processing and bioinformatics

DIA raw files were analyzed using Spectronaut (version 20.1, Biognosys) with the built-in Pulsar search engine against the Uniprot *Homo sapiens* reference proteome (#UP000005640, downloaded on 11th September 2025). Search parameters included full trypsin specificity, 7–52 amino acid length, and a maximum of two missed cleavages. Carbamidomethylation at cysteine was used as a fixed modification, and protein N-terminal acetylation and methionine oxidation were set as variable modifications. The false discovery rates (FDRs) were set as 0.01 for the peptide-spectrum match (PSM), peptide, and protein identification. For quantification, the identified (Qvalue) was set for precursor filtering and MS2 quantification with the area as quantity type. The original mass spectrometric raw Spectronaut files are available on the proteomeXchange PRIDE platform^63^.

#### Proteomics analysis

For protein quantification, protein intensities ≤ 1 were set to missing (NA) on a per-sample basis. Protein groups were then filtered based on data completeness within genome-defined sample groups: protein groups were retained if they were detected (non-NA) in at least 50% of samples in either the ROY or the LOY genome group.

Differential expression analysis was performed independently for each condition. For each comparison, the NA-filtered expression matrix was subset to samples corresponding to the given condition. Linear modeling was carried out using the limma package in R^64^ to assess differences between genome groups (LOY vs. ROY). A design matrix of the form Intensity ∼ Genome was fitted, and differential expression was estimated for LOY relative to ROY. Empirical Bayes moderation (eBayes) was applied, and moderated t-statistics were obtained with false discovery rate (FDR) correction. Further analysis and graphical representation were conducted within the R framework^65^, incorporating, among other packages, tidyverse^66^ for data manipulation and visualization.

Functional enrichment analysis was performed using gene set enrichment analysis (GSEA) as implemented in the clusterProfiler R package^67^. For each analysis, proteins were ranked according to their log2 fold change derived from the differential expression results. Enrichment was performed against the Hallmark gene set collection from the Molecular Signatures Database (MSigDB)^68^, using as background universe all proteins detected in the filtered proteome. Gene sets containing fewer than 5 or more than 500 members were excluded.

### Immunofluorescence (IF) staining

60,000 cells were seeded per well in 96-well glass-bottom plates (Greiner). After 24 h, cells were washed with 1X PBS, MgCl_2_, and CaCl_2_ (Gibco), fixed with 10% formalin (Sigma-Aldrich) for 10 min, washed, and permeabilized with 0.1% Triton X-100 (Sigma-Aldrich) for 10 min at room temperature and washed again. Non-specific binding was blocked by incubation in the blocking buffer (5% BSA, 2% Goat Serum in PBS) for 1 h at room temperature. All antibody dilutions were done in blocking buffer. Cells were incubated with primary antibodies against cadherins (rabbit anti–N-cadherin (1:100, Cell Signaling Technologies) or rabbit anti-E-cadherin (1:100, Cell Signaling Technologies)) overnight at 4 °C.

Cells were washed three times with PBS (Gibco) and incubated with goat anti-rabbit IgG AF488 (1:500, Invitrogen) for 1 h at room temperature in the dark. Cells were washed three times with PBS (Gibco), counterstained with DAPI (1:1000, Millipore) for 15 min at room temperature, washed again, and stored in PBS at 4 °C until imaging. Images were acquired at the Light Microscopy Imaging Facility of the German Cancer Research Center (DKFZ) using a Zeiss LSM 980 Airyscan confocal microscope. Images were processed using the ZEN2 software (blue edition) (Carl Zeiss).

### CD90 Flow cytometry

For CD90 analysis, 350,000 cells were seeded per 10-cm dish and cultured for 2 days, after which 800,000 cells were harvested. Cells were detached using TrypLE Express (Gibco) for 5 min, washed, and resuspended in a wash buffer (PBS supplemented with 2% FBS). Cells were stained with anti-CD90 antibody (1:50, Miltenyi Biotec) or the corresponding isotype control (1:50, Miltenyi Biotec) diluted in wash buffer for 30 min at room temperature in the dark. Cells were washed twice, resuspended in wash buffer, and transferred to flow cytometry tubes. Unstained controls were prepared from pooled aliquots of each clone. Data were acquired on a BD FACSCanto II flow cytometer at the Flow Cytometry Core Facility, German Cancer Research Center (DKFZ), and analyzed using Kaluza software (v1.3.1; Beckman Coulter).

### DiMeLo-seq

DiMeLo-seq was performed according to published protocols²⁹˒³⁰, with minor modifications.

#### Buffer and Complex Preparation

50mL of wash buffer (20 mM HEPES-KOH, 150 mM NaCl, 0.5 mM spermidine, 0.1% BSA, and EDTA-free Complete™ protease inhibitor) was freshly prepared and filtered (0.2 µm). This buffer was supplemented with either 0.02% digitonin (Digitonin Buffer) or 0.1% Tween-20 (Tween Buffer). To generate the targeting complexes, pA-Hia5 (200 nM) was incubated with H3K4me3 antibody (1:50; Diagenode) in Tween Buffer for 40 min at room temperature.

#### Nuclei Isolation and H3K4me3 Targeting

Cryopreserved cells (3 × 10⁶) were thawed, washed with PBS, and pelleted using a swinging-bucket rotor centrifuge (500 x g, 3 min, 4°C). Cells were resuspended in Digitonin Buffer and incubated on ice for 5 min. Nuclei isolation efficiency was verified via NucleoCounter (Solution 18), with only samples showing <5% viability (indicating successful permeabilization) proceeding to downstream steps.

Isolated nuclei (3 × 10⁶) were pelleted, resuspended in the preformed pA-Hia5–antibody complex, and incubated on a rotator at 4°C for 2 h. Nuclei were then washed twice with Tween Buffer (500 x g, 3 min, 4°C) to remove unbound complexes.

#### Targeted Methylation and Nuclei Lysis

To activate Hia5 methylation in the proximity of H3K4me3 modifications, nuclei were resuspended in Activation Buffer (15 mM Tris-HCl pH 8.0, 15 mM NaCl, 60 mM KCl, 1 mM EDTA, 0.5 mM EGTA, 0.05 mM spermidine, 0.1% BSA) supplemented with 800 µM S-adenosylmethionine (SAM). Samples were incubated at 37°C for 2 h, with gentle tapping every 30 min, and a freshly prepared SAM spike-in (2.5 µL) was added after 60 min. Following activation, nuclei were pelleted, resuspended in PBS, and lysed using Proteinase K and Buffer AL (Qiagen) at 56°C for 10 min. Lysates were stored at 4°C overnight before DNA isolation.

#### DNA extraction and quality control

DNA was purified using the QIAGEN DNeasy Blood & Tissue Kit, starting from the ethanol addition step. Briefly, 200 µL of 100% ethanol was added to each lysate and mixed thoroughly by vortexing. The mixture was transferred to a DNeasy Mini spin column and following manufacturer’s instructions. In the final step, DNA was eluted twice with 75 µL Buffer AE. The DNA concentration and fragment size were determined using the Qubit™ dsDNA BR Assaykit (Invitrogen) and the Agilent TapeStation Genomic DNA ScreenTape, respectively. Size selection was performed using PacBio SRE XS kit, following the manufacturer’s protocol without modification. After size selection, the DNA concentration and fragment size were reassessed using Qubit (Invitrogen) and TapeStation (Agilent).

#### Library preparation and sequencing

Nanopore sequencing was performed on PromethION flow cells (FLO-PRO114M, R10.4.1; Oxford Nanopore Technologies) using the Oxford Nanopore Technologies Ligation Sequencing Kit v14 (SQK-LSK114) and the protocol *Human variation sequencing from 30 kb extracted cell line samples using SQK-LSK114* (GDH_9174_v114_revC_10Nov2022-promethion). Flow cells were washed when the number of active pores dropped below 20% using the *Library Recovery from Flow Cells* protocol (LIR_9178_v1_revK_13Dec2024), following the “Recover a library to replace on a washed flow cell” section. When an additional library was available, it was loaded directly instead of replacing the recovered library.

#### Basecalling and alignment

Raw nanopore sequencing data in POD5 format were base-called using Dorado v1.1.0 with the DNA_R10.4.1_E8.2_400bps_HAC@v5.2.0 model with simultaneous alignment to the human reference genome (hg38). DNA base modifications were detected during basecalling using the corresponding Dorado modified-base models for adenine methylation (N6-methyladenine; 6mA) and CpG-associated cytosine modifications (5-methylcytosine and 5-hydroxymethylcytosine; 5mC/5hmC). Adenine methylation (6mA) was used as a proxy for protein–DNA interactions, reflecting paHia5-mediated methylation at genomic loci bound by H3K4me3-associated chromatin. CpG-associated cytosine modifications were analyzed in parallel to assess endogenous DNA methylation patterns. The resulting BAM files were processed using samtools (v1.20) for sorting and indexing, and bamtools (v2.5.2) for downstream BAM file manipulation and quality control before analysis. DNA modification calling and aggregation were performed using ModKit (v0.3.1; Oxford Nanopore Technologies) pileup, which extracts per-base DNA modification probabilities from aligned reads. For each aligned BAM file, two bedMethyl files were generated: one reporting CpG-associated cytosine modifications (5mC/5hmC) and one reporting adenine modifications (6mA). To ensure high-confidence modification calls, only bases with a minimum modification probability of ≥ 0.9 were retained for downstream analyses. Bedtools (v2.31.1) was used for genomic interval operations on bedMethyl files, including intersection with annotated genomic features.

#### CpG island–level DNA methylation analysis

CpG island–level DNA methylation analysis was performed using RnBeads (v2.24.0) in R (v4.4.3). Preprocessed methylation data from eight A549 clones (four LOY and four ROY) were imported as *bedmethyls* files. Differential methylation between LOY and ROY groups was assessed at the CpG island level, producing mean methylation differences and combined rank statistics. CpG islands on sex chromosomes were excluded, and remaining islands were assigned to the nearest transcription start site using Ensembl annotations (EnsDb.Hsapiens.v86, hg38); islands located more than 10 kb from any TSS were discarded. Non-coding genes (antisense transcripts, lncRNAs, miRNAs, rRNA genes, and RP-prefixed pseudogenes) were filtered out. For genes mapping to multiple CpG islands, the island with the largest absolute mean methylation difference was retained, and differentially methylated islands were defined using an absolute mean methylation difference threshold of 0.2. Mean CpG island methylation values were visualized using (ComplexHeatmap; v2.22.0) with samples ordered by group (LOY then ROY). Group-level methylation distributions were compared using boxplots (ggplot2; v4.0.1).

#### DMR calling

DMRs between LOY and ROY A549 clones were identified using the DSS Bioconductor package (v2.48.0) in R (v4.3.3). CpG-level methylation data were imported from *bedMethyl* files using bsseq (v1.46.0) via the read.bedMethyl() function, generating a BSseq object with strand-collapsed CpGs, zero-coverage sites removed. CpG-level differential methylation was assessed using a Wald test under a beta-binomial regression model accounting for biological variability across replicates. Prior to testing, methylation levels were smoothed using a 500 bp moving-average window. Differentially methylated CpGs were defined using a p-value threshold of 0.05 and a minimum absolute methylation difference (Δβ) of 0.2. Significant CpGs were merged into DMRs using callDMR() with the same thresholds, requiring a minimum of three CpGs per region, a minimum region length of 50 bp, and a consistent direction of methylation change across all CpGs within each region.

#### DMR annotation, pathway enrichment, and locus-specific methylation visualization

DMRs were annotated to genomic features and associated genes using ChIPseeker (v1.42.1) in R (v4.3.2). DMRs located on the Y chromosome were excluded from downstream functional analyses. For functional interpretation, GSEA was performed using a preranked approach implemented in GSEApy (v1.1.11; Python v3.10.19). Genes were ranked based on promoter-associated DNA methylation changes derived from DMRs overlapping promoter regions (±5 kb from transcription start sites). For genes associated with multiple promoter-overlapping DMRs, a single representative score was assigned by selecting the DMR with the maximum absolute test statistic (areaStat), retaining its original sign. To facilitate biological interpretation, the sign of the areaStat was inverted before ranking, such that positive values indicate promoter hypomethylation and negative values indicate promoter hypermethylation. Pre-ranked GSEA was conducted using the MSigDB Hallmark gene set collection (v7.1). Gene sets containing fewer than 10 or more than 500 genes were excluded. Enrichment significance was assessed using 1,000 permutations with a fixed random seed (seed = 6) to ensure reproducibility. Resulting p-values were adjusted for multiple testing using the false discovery rate (FDR). Single-molecule CpG methylation profiles at selected gene loci were visualized using the NanoMethViz R/Bioconductor package (v2.8.1). Eight A549 cell line clones were grouped according to Y chromosome status and modified base BAM files were imported directly using the ModBamFiles interface. Gene annotations were obtained from the Ensembl database (EnsDb v86; hg38) and restricted to canonical chromosomes (chr1–22, chrX, chrY). CpG sites with fewer than 10 supporting reads were excluded (NanoMethViz.site_filter = 10). Gene-level methylation was visualized using plot_gene(), which generates group-wise smoothed methylation probability profiles (LOESS smoothing, span = 0.05) along with gene model tracks.

#### Identification of H3K4me3 signal

Peak assessment was performed using a method inspired by the identification of accessible chromatin in Lee et al^33^. First, the adenine methylation frequencies were averaged across non-overlapping 10-bp bins along each chromosome. Regions in which the average adenine methylation frequency exceeded the 99th percentile of the chromosome’s distribution were selected as candidate regions. The significance of each region was assessed using a binomial test of the raw methylation frequency, with the overall methylation frequency of the respective chromosome serving as the null probability. P-values were adjusted for multiple testing using the Benjamini–Hochberg correction. Regions with an adjusted p-value of less than 0.01 and a minimum width of 100 bp were designated as significant peaks. This procedure was applied to all chromosomes independently to identify peaks across the genome using R (4.4.2). To avoid having multiple peaks in nearby regions, peaks within 1kB distance of one another were merged. The resulting peaks were annotated to genomic features using the ChIPseeker (1.42.1).

#### Differential analysis of H3K4me3 signal

Differential analysis of adenine-methylation was performed by extracting adenine-methylation calls (6mA) in regions 2 kb up- or downstream of TSS using bedtools (v2.31.1). In R (4.4.3), the following steps were conducted to calculate log2FC between LOY and ROY clones. For each sample, the calls were filtered for positions that had at least 10x sequencing depth. To account for global differences in adenine-methylation that might be technical artifacts, the mean methylation fraction per sample and among all samples was calculated. We then computed sample-specific scaling factors by dividing the mean of all samples by the sample-specific mean and multiplied the methylation levels by this scaling factor. log2FC was calculated between the average for LOY and ROY clones for each region. These values were used to rank genes for GSEA, with GSEApy (v1.1.11; Python v3.10.19). Enrichment was assessed using the MSigDB Hallmark gene set collection (v7.1). Enrichment significance was determined using 1,000 gene set permutations, and enrichment results were corrected for multiple testing using the false discovery rate. For visualization of adenine-methylation intensity around regions of interest, methylation fractions were calculated in 20 bp bins throughout the genome using Python (3.11.9). The scaling factors calculated from the mean adenine-methylation in promoter regions were applied to these tracks, and the averages across all LOY clones and ROY clones were computed. Scaling and averaging were performed using deeptools (v3.5.6)^69^. Then, the tracks were used for visualization in regions of interest using pygenometracks (v3.9)^70^. H3K4me3-associated 6mA signal from DiMeLo-seq was extracted from bigWig files over differentially methylated CpG islands (CGIs) for each clone. Mean 6mA signal per CGI was calculated, Z-score normalized across clones, and visualized as a heatmap.

### Cytometry by Time of Flight (CyTOF)

6 million cells were harvested and washed with Maxpar CyTOF PBS (Fluidigm). Live/dead staining was conducted by resuspension of the cells with 1.25µM Cisplatin (Fluidigm) in CyTOF PBS for 1 minute. Cisplatin was neutralized with medium supplied with 10% FBS (Invitrogen). Fixation and permeabilization occurred in two steps: first, cells were incubated in 1.6% PFA, then washed with Maxpar Cell Staining Buffer (Fluidigm). Second fixation and permeabilization was conducted by incubation of cells with Maxpar nuclear antigen staining buffer set (Fluidigm) according to the manufacturer’s instructions. Samples were multiplexed by taking 3 million cells/sample and using the Cell-ID 20-Plex Pd Barcoding Kit (Fluidigm). After barcoding, 0.5 million cells from each sample were pooled to prevent batch effect and blocked using FBS for 10 minutes. Then, the pooled sample was directly incubated for 30 minutes at room temperature with the mixture of antibodies and washed with Maxpar Cell Staining Buffer. Cells were fixed with fresh 4% PFA overnight. On acquisition day, samples were suspended with 4% PFA supplied with Cell-ID Intercalator-Iridium (Fluidigm) at 125nM. Cells were then washed with Maxpar Cell acquisition solution (Fluidigm) and suspended again in Maxpar Cell acquisition solution supplied with 10% EQ Four Element Calibration Beads (Fluidigm). Before the run on the CyTOF machine, cells were filtered through a 35µM mesh cell strainer. Data was acquired on a Fluidigm Helios CyTOF system. The full panel is provided in **Table 13**. Antibodies (BSA and azide-free) were conjugated to metals using the MIBItag Conjugation Kit (IONpath) according to the manufacturer’s instructions.

#### Data analysis

CyTOF data underwent the following pre-processing before analyses: First, the CyTOF software by Fluidigm was used for normalization and concatenation of the acquired data. Then, several gates were applied using the Cytobank platform (Beckman Coulter):

The normalization beads were gated out using the 140Ce channel. Then, live single cells were gated using the cisplatin 195Pt, iridium DNA label in 193Ir, event length, and the Gaussian parameters of width, center, offset, and residual channels. CyTOF software was then used for sample de-barcoding.

Next, the downstream pipeline followed our work^34,36^, with some modifications.

Cells were gated using the core histones (H3, H3.3, and H4), and only cells with a minimum raw value of 5 for all core histones were included. For the various markers, a hyperbolic arcsine transformation (with a scale factor of 5) was first applied to the data:

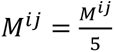

The data were normalized using the measured values of the core histones:

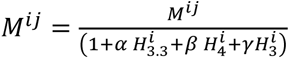

As in^71^, the coefficients were selected to minimize the sum of the variances of the core histones, with *γ* = (1 − *⍺* − *β*). A reduction of a few dozen percent in the sum of variances was observed after the subtraction, indicating the removal of systematic effects.

Finally, the data were z-transformed:

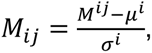

where i denotes the observation, j denotes the column, *μ*^*i*^ and *σ*^*i*^ are the column mean and standard deviation, respectively. Z-transformation was performed in parallel on samples acquired together.

For heterogeneity analysis, we calculated the nearest-neighbor distance (NND), which refers to the distance between each point and its nearest neighbor. We then plotted the ratio of the cumulative distribution functions (CDFs) of LOY cells versus ROY cells.

### Single-cell Multiome

Nuclei Isolation was performed according to 10X Genomics demonstrated protocol (Nuclei Isolation for Single Cell Multiome ATAC + Gene Expression Sequencing - CG000365) starting with 5×10^5^ fresh cells. Cells were incubated with the lysis buffer for 4 minutes. Single-nuclei ATAC and 3’ gene expression sequencing libraries were generated using the Chromium Next GEM Single Cell Multiome ATAC + Gene Expression Reagent Bundle (10X Genomics) according to the manufacturer’s instructions. Per well of the Chromium Next GEM Chip J (10X Genomics) one sample with 15,000 nuclei was loaded. Libraries were sequenced on a NovaSeq 6000 system using the S4 v1.5 flow cell with paired-end 200 cycles (1 lane for gene expression and 3 lanes for ATAC libraries).

Raw read data were processed using 10x Genomics’ Cellranger-ARC v2.0.2 software using default parameters to generate gene count matrices and fragment files for snRNA-seq and scATAC-seq, respectively. We analyzed snRNA-seq and scATAC-seq data separately. For the snRNA-seq data, we filtered cells with a mitochondrial content 3-times higher than the median of all cells, and with less than 5-times the number of features. Doublets were removed with scDblFinder^72^. Low-dimensional representations were generated using standard Seurat v5.3.0 processing^73^. For scATAC-seq data, downstream analyses were performed with R v4.4.3 using the ArchR R package v1.0.3^74^. Cells with TSS enrichment scores less than four or with more than 1000 fragments were removed, and doublets were filtered out using default parameters. Dimensionality reduction was performed via iterative Latent Semantic Indexing (LSI), followed by clustering using the Seurat v5.3.0 method. The EMT scores were computed using Seurat’s function AddModuleScore using the gene set described in **Table 9**. For the scATAC-data, gene score matrices were generated with addGeneScoreMatrix() function to assess accessibility at promoter regions (2.5 kb upstream and 2.5 kb downstream of the transcription start site) for EMT genes. Accessibility tracks were visualized using the plotBrowserTrack() function.

### *In vivo* animal studies

A549 tail vein injection experiments were performed in accordance with and approved by the Landesamt für Natur, Umwelt und Verbraucherschutz Nordrhein-Westfalen, Germany (LAVE, former LANUV). Immunodeficient NOD/Scid/ᵧC (NSG) mice were originally obtained from The Jackson Laboratory via Charles River (Stock Number #005557). Animals were kept in stable groups of 2-5 animals in individually ventilated cages with standard bedding. Water and standard laboratory mouse diet were provided ad libitum. Water was filtered into autoclaved bottles. Bedding was regularly changed under a hood to preserve sterile conditions. Light/dark rhythm was set at 12:12 hours, and room temperature and humidity were kept stable.

For the experiment, both male and female adult mice from the age of 8 weeks or older were equally used. All animal experiments and data analyses were performed without blinding, and no power analysis was conducted to pre-determine group size. All animals were tagged with ear marks for identification before the experiments. No further procedures were performed besides the described experiments in this manuscript.

To assess survival in the bloodstream, seeding and outgrowths in the lung, 1 × 10^5^ A549 cells were injected intravenously into the lateral tail vein of NOD/Scid/ᵧC (NSG) mice. Six weeks after injection, mice were sacrificed, and lung lobes were preserved for histological analysis as FFPE tissue and used for dual CISH staining. As for control, a cyto-spin pellet of the injected A549 cells (∼1×10^6^ cells) was paraffin-embedded and prepared for CISH analysis.

### Chromogenic in situ Hybridization (CISH) and LOY/ROY quantification

Dual-color Chromogenic in situ Hybridization (CISH) was performed to detect X and Y chromosomes in FFPE tissue sections. We utilized the ZytoDot® 2C CEN X/Y Probe (ZYTOMICS), targeting centromeric regions, in combination with the ZytoDot® 2C CISH Implementation Kit (ZYTOMICS). All steps, including pretreatment, denaturation, hybridization, and detection, were performed according to the manufacturer’s instructions.

Stained slides were digitized using a 3DHISTECH Pannoramic digital slide scanner at 40× magnification. Digital image analysis was performed using QuPath (v0.6.0) open-source software^75^. For quantification, individual tumor nodules were annotated and inspected at high resolution to classify cells based on the presence of Y-chromosome (green) and X-chromosome (red) nuclear signals.

Tumor nodules were categorized based on the Y-signal as follows:

LOY tumors: nodules containing >90% LOY cells (loss of Y signal).

ROY tumors: nodules containing >80% ROY cells (retention of dual X/Y signals).

Mixed tumors: nodules containing heterogeneous populations of both LOY and ROY tumor cells.

To ensure robustness, scoring was blinded and quantified by two independent researchers, with histological review performed by a certified pathologist. Data are reported as the percentage of tumor nodules per category per animal.

### Single-cell RNA-seq and lineage tracing data from published metastatic A549 mouse xenograft models

A549 single-cell RNA-seq datasets from Quinn *et al*.^39^ were reanalyzed using Python (v3.10.12) and the Scanpy framework (v1.11.0), focusing on the pre-implantation cells (LM0; collected *in vitro* before xenografting) and mouse M5k (established by injecting 5,000 engineered A549-lineage-tracing cells directly into the left lung). Standard quality control analysis was conducted. Cells with 200-7,000 detected genes and less than 10% mitochondrial reads were retained, and genes detected in fewer than three cells were excluded. Read counts were normalized and log-transformed with one pseudocount addition. Highly variable genes were selected (min_mean=0.0125, max_mean=3, min_disp=0.5). Principal component analysis (100 components) was followed by batch correction using Harmony (theta = 2, max_iter = 100). Neighborhood graphs were constructed using the top 40 principal components (n_neighbors = 15), UMAP embeddings were generated (min_dist = 0.5, spread = 1.0). Sex-chromosome and housekeeping gene signatures were scored using Scanpy’s score_genes function, requiring at least three signature genes per cell and 50 control genes, yielding Y_score, X_score, and Houskeep_score. Y-score distributions were assessed by Gaussian mixture modeling, and cells were classified as LOY (Y-score < −0.2) or ROY. Y-score associations with metastatic capacity (scMetRate, derived from scTreeMetRate in the cellmeta data provided by the original study) were evaluated using Spearman correlation. For clone-level analysis (Figure S6H), clones (LineageGroup) with ≥20 cells per site were classified as LOY- or ROY-enriched if the corresponding fraction was ≥0.75 across sites (mixed excluded). Clone metastatic phenotype was defined by the median clone TreeMetRate (any-metastatic vs non-metastatic), and differences between LOY and ROY clones were tested using Fisher’s exact test on clone-count contingency tables.

## Data and code availability

The mass spectrometry proteomics data have been deposited to the ProteomeXchange Consortium via the PRIDE^76^ partner repository with the dataset identifier PXD074050. Processed CyTOF data is available from figshare (doi provided after publication, reviewer access: https://figshare.com/s/8c1c06c6c060bd32987c). Single-cell RNA sequencing data from the lung donors are available as processed Seurat objects from figshare (https://figshare.com/s/337294e217b299c3f2a6). A549 multi-omic data is available from https://figshare.com/s/0bb9ea2ee2e0ea53f74c. The raw data generated in this study are available via controlled access in the German Human Genome-Phenome Archive under the GHGA Accession Number https://data.ghga.de/study/GHGAS27424019412227 (will become public upon publication). Further details, including the data access policy for the study, can be found there. Code to reproduce the figures of the manuscript is available from: https://git.dkfz.de/b370/lung_group/loy_cells.

## Declaration of generative AI and AI-assisted technologies in the manuscript preparation process

During the preparation of this work, the author(s) used Google Gemini to consolidate language across authors and sections, remove repetitive statements, and increase clarity. Claude and ChatGPT were used to troubleshoot coding errors and installation issues. After using these tools, the authors reviewed and edited the content as needed and take full responsibility for the content of the published article.

## Acknowledgement

The authors would like to thank the following core facilities at DKFZ: Genomics and Proteomics, Flow Cytometry, scOpenLab, Light Microscopy, Open Sequencing Lab, NGS, and the DKFZ Omics IT and Data Management for their excellent support. We thank Aaron Straight for providing us with the plasmid pET-pA-Hia5 (Addgene, 174372) and the EMBL Protein Expression and Purification Core Facility for the production of pA-Hia5, and Alex Vogel (Oxford Nanopore Technologies) for technical support and guidance. We are grateful to Christa Stolp, Martin Fallenbüchel, and Maja Gräbner (Thoraxklinik Heidelberg), Alberto Diaz-Jimenez (DKFZ), Barbara Helm (DKFZ), and Agustin Rodriguez (DKFZ) for their support with human sample collection. We thank Yaara Oren for great input and discussions. We would also like to acknowledge the excellent technical support of Marion Bähr and Jessica Heilmann, and members of the Division of Cancer Epigenomics for helpful discussions. Most importantly, we thank the patients and their families for their generosity in donating samples to this study.

Michael Scherer is supported by a postdoctoral fellowship from the Dr. Rurainski Foundation. Mei-Ju May Chen and Dung-Chi Wu are supported by the Yushan Fellow Program (Ministry of Education, MOE, Taiwan). This work was supported in part by the Helmholtz International Graduate School (K.S., G.A.), the DFG CRC 1709 project A04 (C.P., M.L.P.), the DFG CRC 1366/2 project A5 (C.P.), the Wilhelm Sander-Stiftung project 2024.117.1 (M.S.), the German Center for Lung Research (DZL), funding codes 82DZL004C4 and 82DZLT84C4 [C.P., M.L.P., R. So., U. K.], 82DZL004C2 and 82DZLT84C2 [H.W, L.V.K., F.J.F.H, M.A.S.], 82DZL004C1 [D.K., A. St.], 82DZL005C3 [R. Sa.], the German Cancer Aid project 70114056 (P.M., B.M.G.), and an Add-on Fellowship from the Joachim Herz Stiftung (K.S.).

## AUTHOR CONTRIBUTIONS

Conceptualization: K.S., M-J.M.C., M.S., C.P., M.L-P.

Methodology: K.S., M-J.M.C., R. M., D. K., A. St., U.K., B. M. G., G.R., E. S., M.S., C.P., M.L-P

Software: K.S., M-J.M.C., G. A., S. M-S., D-C. W., N. Z., L. H., N. E., K. K., E. S., A. F-S., P. L., G. R., M. S.

Formal analysis: K.S., M-J.M.C., G. A., S. M-S., D-C. W., N. Z., L. H., N. E., A. F-S, D. K., A. St., M. Schi., G. R., E. S., M.S., M.L-P.

Investigation: K. S., M.-J. M. C., G. A., S. M.-S., D.-C. W., N. Z., O. G., L. H., F. B., J. C., N. E., O. M., R. M., K. K., S. C., E. S., A. F.-S., C. S., M. J. A.-D. G., S. M., M. D., P. M.

Visualization: K. S., M.-J. M. C., G. A., S. M.-S., D.-C. W., L. H., N. E., A. F-S., G. R., M.S., M.L-P.

Validation: M-J.M.C., M.S., C.P., M.L-P.

Data Curation: K.S., M-J.M.C., M.S.

Resources: K. K., E. S., M. D., P. M., B. H., H. W., L. V. K., M. K., F. J. F. H., T. M., M. A. S., A. S.

Writing – original draft: K.S., M-J.M.C., M.S. C. P., M. L-P. Writing – review & editing: all authors

Supervision: F. J. H., P. L., M. Schi., U. K., Ro. So., R. S., B. M. G., G. R., E. S., M. S., C. P., M. L.-P.

Project administration: M.S., C. P., M. L-P.

Funding acquisition: M.S., C. P., M. L-P.

---

**Figure S1:**
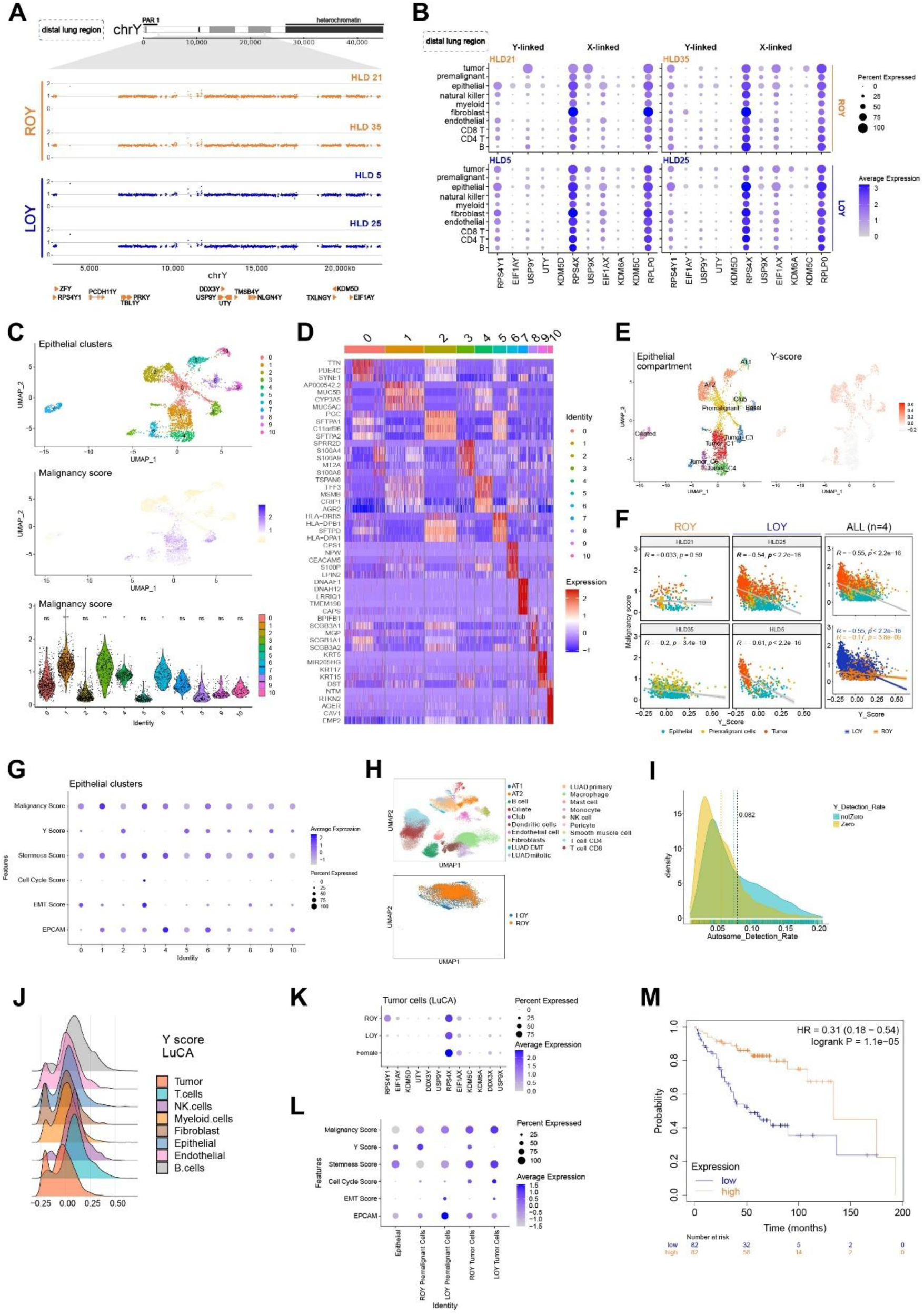
LOY in LUAD is enriched in tumor cells and associates with adverse clinical outcomes, related to Figure 1. **(A)** WGS profiles of distal lung tissue from ROY (n=2) and LOY (n=2) patients, showing a zoomed-in view of the Y chromosome. **(B)** Dot plots showing expression of five Y-linked genes, X-linked homologs, and *RPLP0* in distal lung tissue. Dot size indicates proportion of cells expressing each gene; color intensity represents mean expression. **(C)** UMAP projection of epithelial cells from tumor and distal-lung tissues, colored by cluster (top) and malignancy score (middle). Corresponding violin plots (bottom) displaying the malignancy score per cell. **(D)** Heatmap showing the mean expression of the top five marker genes per epithelial cluster. **(E)** UMAP projections of the epithelial compartment from all regions, colored by annotation (top) and Y-score (bottom). **(F)** Correlation between malignancy score (y axis) and Y-score (x axis) for epithelial, premalignant, and tumor cells for ROY tumors (left), LOY tumors (middle), and for the combined cohort (right). **(G)** Dot plot displaying malignancy, Y, stemness, cell cycle, and EMT scores, alongside *EpCAM* expression across epithelial clusters. **(H)** UMAP projections of all cells from the LuCA dataset^26^ colored by annotation (top), and tumor cells colored by LOY/ROY status (bottom). **(I)** Density plot of autosome detection rate for cells with (notZero, cyan) or without (Zero, yellow) Y-chromosome gene detection. **(J)** Density plots showing Y-score distributions for all annotated cell types in the LuCA dataset^26^. **(K)** Dot plot of six Y-linked genes and their X-linked homologs across ROY and LOY tumor cells, including tumor cells from females from the LuCA dataset^26^ as controls. Dot size indicates the percentage of cells expressing each gene; color intensity represents mean expression. **(L)** Dot plot showing malignancy, Y, stemness, cell cycle, and EMT scores, as well as *EpCAM* expression for epithelial cells, premalignant cells (stratified by ROY/LOY status) in the LuCA dataset^26^. **(M)** Kaplan-Meier survival analysis of 328 male LUAD patients stratified by the mean expression of 13 Y-linked genes (*RPS4Y1*, *ZFY*, *PCGH11Y*, *TBL1Y*, *PRKY*, *USP9Y*, *DDX3Y*, *UTY*, *TMSB4Y*, *NLGN4Y*, *TXLNGY*, *KDM5D*, *EIF1AY*) using kmplot.com. Groups represent the lowest (Q1, blue) and highest (Q4, orange) quartiles. Median OS: 52 vs. 133.5 months; HR = 0.31; 95% CI 0.18–0.54, p = 1.1 × 10⁻^5^.

**Figure S2:**
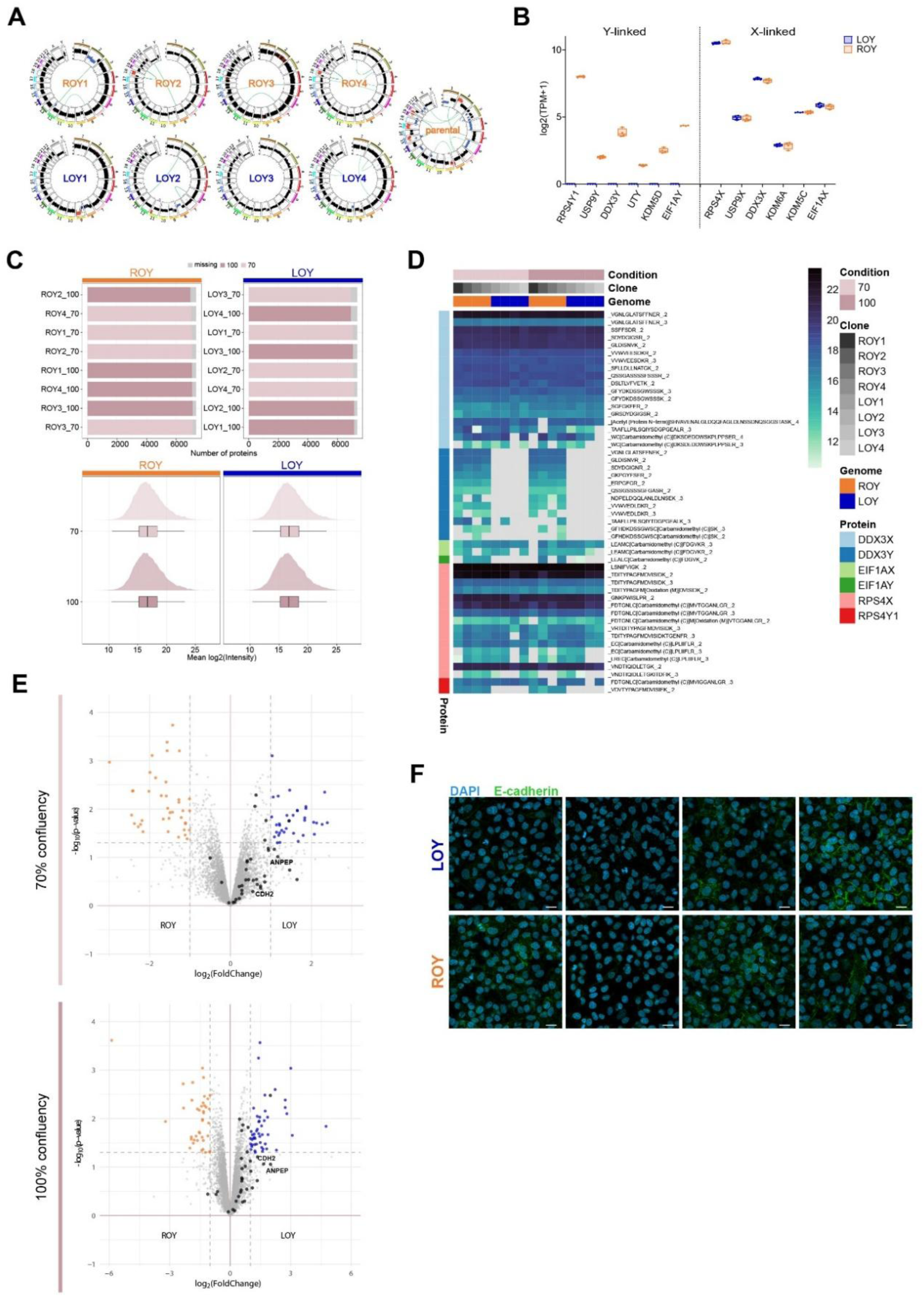
LOY drives intrinsic EMT and inflammatory programs, related to Figure 2. **(A)** WGS profiles of four ROY (top) and four LOY clones (Circos plots) compared to bulk A549 parental cells. **(B)** Boxplot showing expression of six Y-linked genes and X-linked homologs across LOY (n=4, blue) and ROY (n=4, orange) clones (log_2_(TPM+1)). Box represents median and quartiles; whiskers show min/max values. **(D)** Protein detection metrics. Top: proteins detected per sample across confluency conditions. Bottom: mean log2 intensity distribution per group and condition. (**D**) Heatmap of proteotypic peptide intensities for three Y-linked proteins and their X-linked homologs across conditions. (**E**) Volcano plot of differentially abundant proteins at 70% (top) and 100% confluency (bottom). Blue: upregulated in LOY; orange: downregulated in LOY. X-axis: log2 fold change; y-axis: -log_10_(p-value). (**F**) Confocal microscopy images for all eight clones stained for E-cadherin (green) and DAPI (blue). Scale bar: 20 µm.

**Figure S3:**
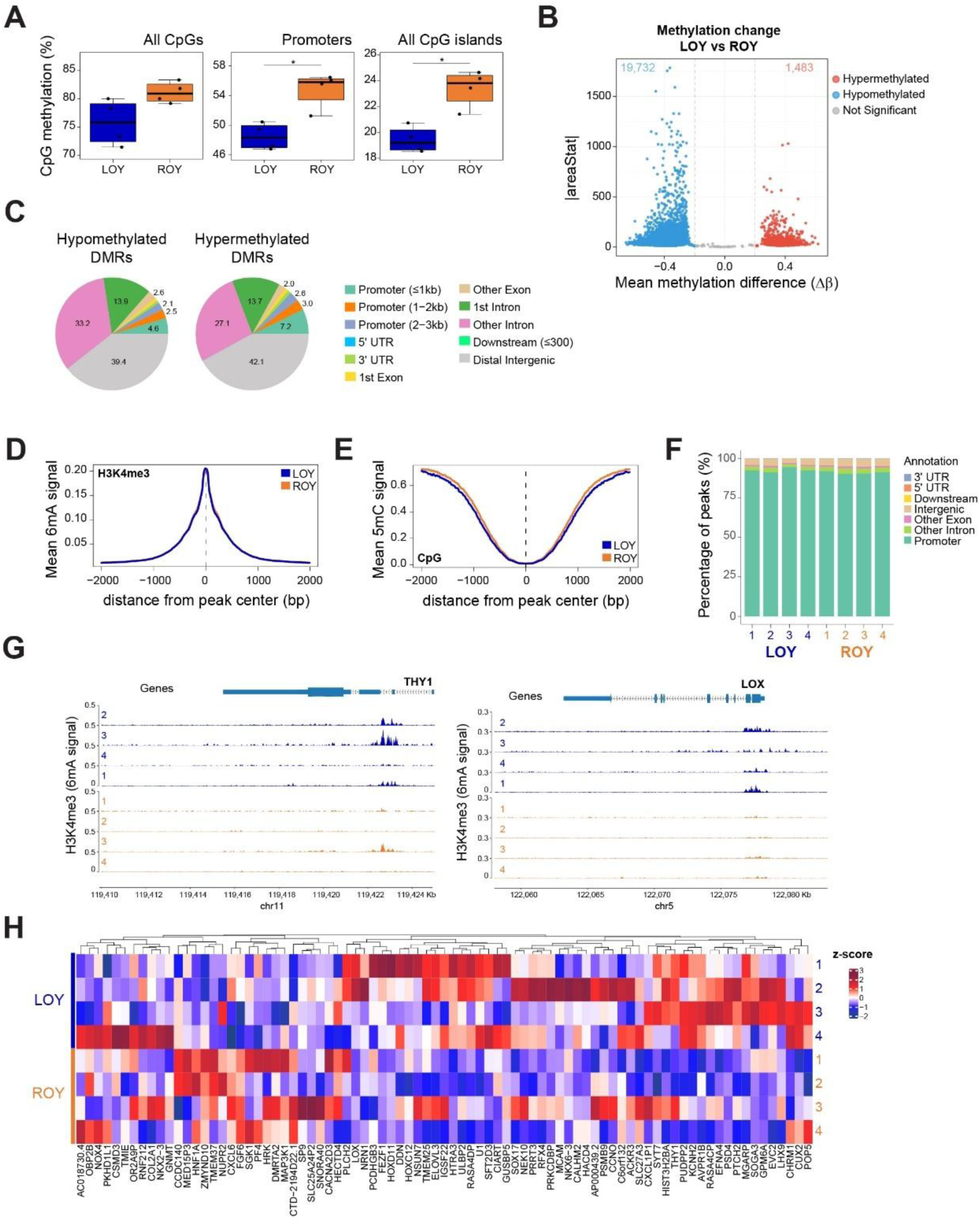
Genome-wide epigenetic alterations in A549 LOY cells reshape the epigenetic landscape through DNA hypomethylation, related to Figure 3. **(A)** Boxplots of mean CpG methylation for all CpGs (left), promoters (middle), and CpG islands (right). Box displays the median and quartiles and whiskers, the min/max values. Statistical significance was assessed by the Wilcoxon rank-sum test (α=0.05). **(B)** Volcano plot of LOY differentially methylated regions (DMRs). X-axis: mean methylation difference (Δβ); y-axis: DSS regional test statistic (|areaStat|). Vertical dashed lines indicate the methylation difference cutoff (|Δβ| > 0.2). **(C)** Genomic annotation of DMRs across genomic features. **(D)** Mean 6mA (H3K4me3) and **(E)** 5mC signal profiles centered on H3K4me3 peaks for ROY (orange) and LOY (blue) clones. X-axis: genomic distance from peak center (0 bp, dashed line); y-axis: mean modification signal. **(F)** Barplots showing genomic annotation of H3K4me3 peaks per clone. **(G)** Locus-specific H3K4me3 signal tracks for *THY1* (left) and *LOX* (right) across all clones. **(H)** Z-score distribution of H3K4me3 signal (6mA) at differentially methylated CpG islands.

**Figure S4:**
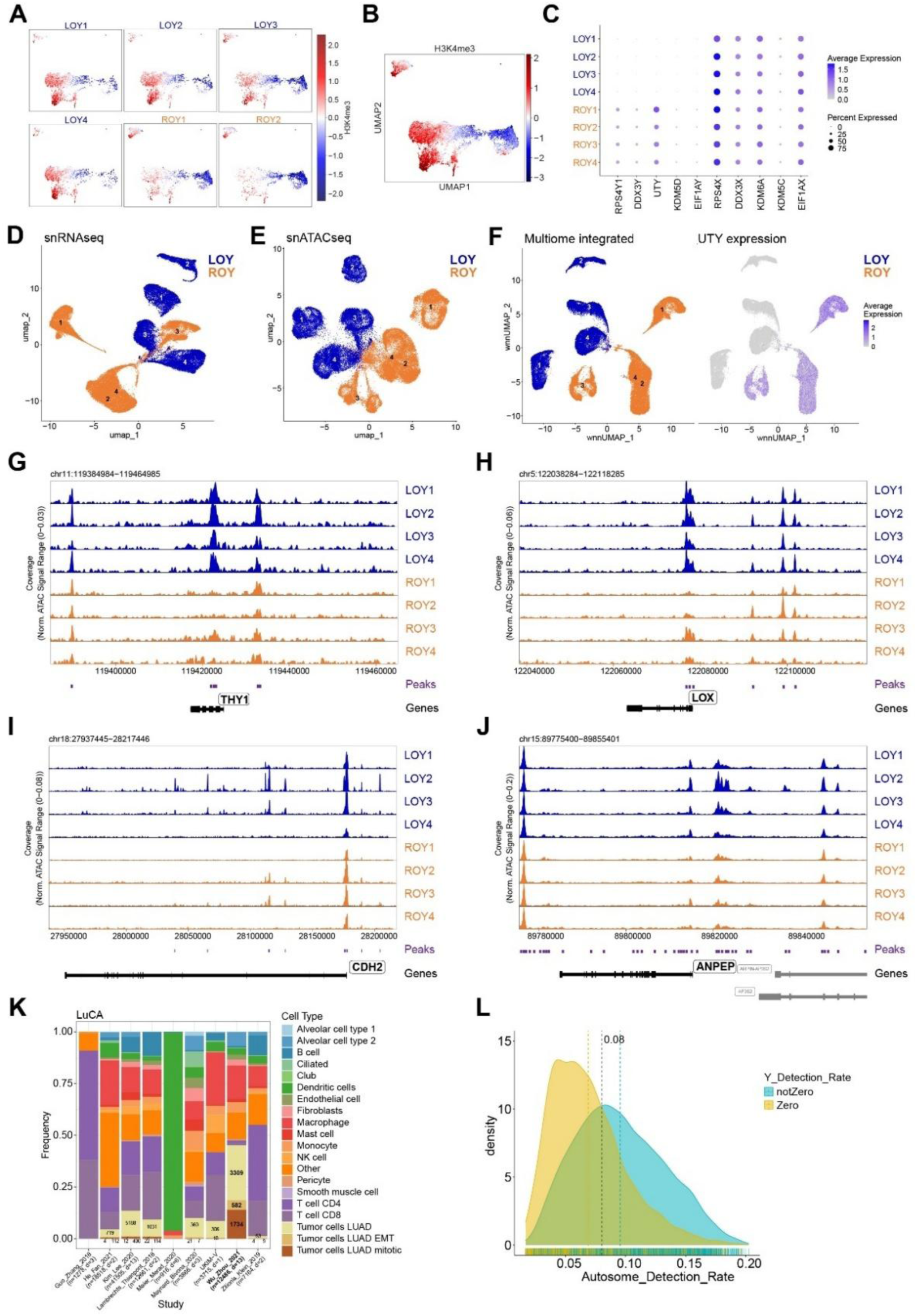
Single-cell profiling reveals LOY-driven epigenetic heterogeneity and transcriptional plasticity, related to Figure 4. **(A)** UMAP projection of Epi-CyTOF data colored by H3K4me3 levels in each sample. **(B)** Integrated UMAP of all samples combined, colored by H3K4me3 expression levels. **(C)** Dot plot of Y- and X-linked gene expression from snRNA-seq across isogenic clones. Dot size indicates the percentage of cells expressing each gene, and color intensity represents the mean expression level. **(D)** UMAP of snRNA-seq data, and **(E)** UMAP of scATAC-seq, colored by LOY/ROY status. **(F)** Multimodal integration UMAP (Seurat’s Weighted Nearest Neighbors, WNN) of snRNA-seq and scATAC-seq data colored by group (left) and *UTY* expression (right). Locus-specific scATAC-seq coverage at *THY1* (G), *LOX* (H), *CDH2* (I), and *ANPEP* (J). **(K)** Stacked bar plot of cell type frequencies per study integrated in the LuCA dataset^26^. Cell numbers are indicated for the three LUAD tumor cell subtypes annotated. **(L)** Density plot of autosome detection rate for cells with (cyan) and without (yellow) Y-gene detection.

**Figure S5:**
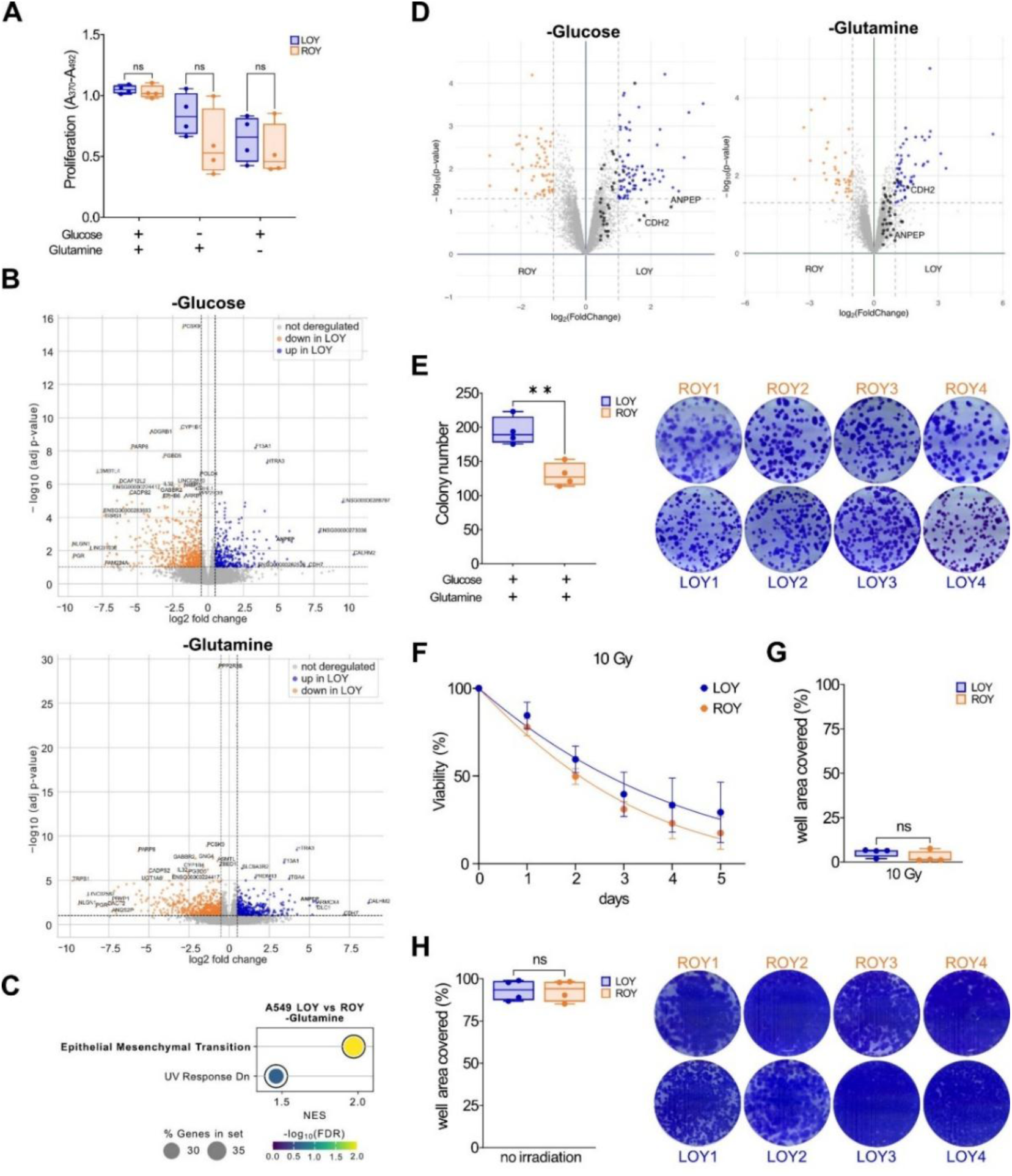
LOY promotes adaptation to metabolic and genotoxic stress, related to Figure 5. **(A)** Boxplots displaying cell proliferation (BrdU ELISA) of four LOY (blue) and four ROY (orange) clones under normal growth conditions (glucose+, glutamine+), glucose deprivation (glucose-, glutamine+), or glutamine deprivation (glucose+, glutamine-). **(B)** Volcano plot of differentially expressed genes between LOY (n=4) and ROY (n=4) clones under glucose deprivation (top) and glutamine deprivation (bottom) (Y-linked genes excluded). X-axis: log2 fold change; y-axis: -log_10_(p-value). Blue: significantly upregulated genes in LOY; orange: downregulated in LOY. **(C)** Dot plot showing the top 2 Hallmark gene sets with FDR<0.25 from preranked GSEA of RNA-seq data comparing A549 LOY and ROY under glutamine deprivation. X-axis: normalized enrichment score (NES); y-axis: gene sets. Dot size indicates gene count; color represents -log_10_(FDR). **(D)** Volcano plots of differentially abundant proteins under glucose (left) and glutamine (right) deprivation. Blue: significantly upregulated in LOY; orange: downregulated in LOY. X-axis: log2 fold change; y-axis: -log_10_(p-value). **(E)** 10-day colony formation assay of four LOY (blue) and four ROY (orange) clones, standard growth conditions. Left: boxplot of mean colony number (n=3 technical replicates/clone). Right: representative images. **(F)** Normalized growth curves of four LOY (blue) and four ROY (orange) clones over 5 days following 10 Gy irradiation, fitted using non-linear regression. **(G)** Boxplot quantifying the area occupied by colonies after 10 Gy irradiation (experiment 1). **(H)** Boxplot quantifying the area occupied by non-irradiated controls (left) and representative images (right) from experiment 1. Box plots as in Figure 5.

**Figure S6:**
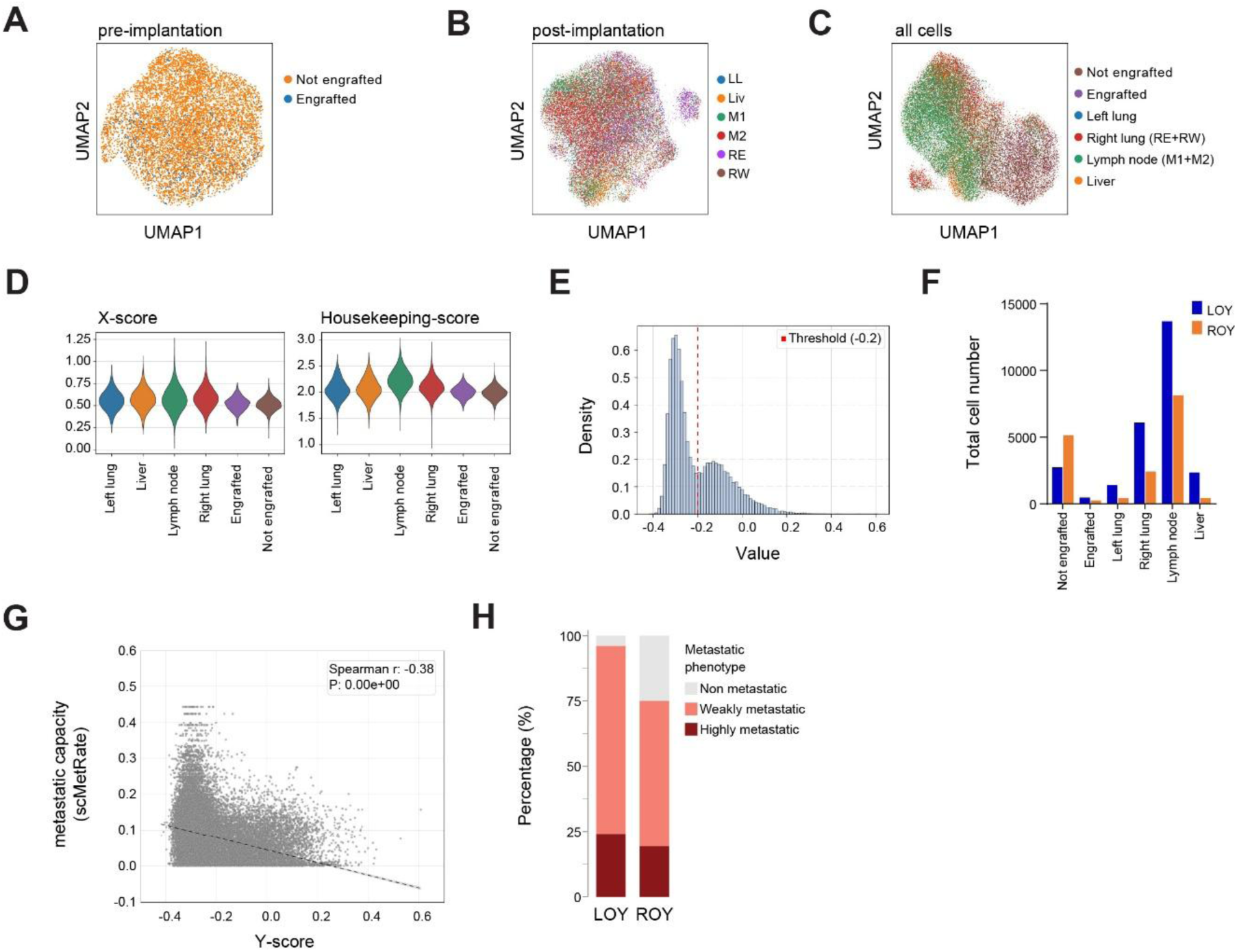
LOY facilitates engraftment and metastatic dissemination, related to Figure 6. All data reanalyzed from Quinn *et al*.^39^ (**A**) UMAP projection of pre-implantation cells (LM0) colored by subsequent engraftment status. **(B)** UMAP projection of engrafted cells (M5K) colored by tissue recovery site. LL: left lung; Liv: liver; M1: mediastinum lymph tissue 1; M2: mediastinum lymph tissue 2; RE: right lung east; RW: right lung west. **(C)** Combined UMAP projection of all cells (pre- and post-implantation) colored by engraftment status and metastasis location. **(D)** Violin plots showing X-score (left) and housekeeping score (right) distributions for each cell across engraftment and metastatic sites. **(E)** Histogram of Y-score distribution showing the classification threshold (red dashed line). (**F**) Bar plot showing the absolute number of cells classified as LOY or ROY per site. (**G**) Spearman correlation between Y-score and scMetRate, colored by metastatic sites. (**H**) Stacked bar plot showing the metastatic phenotype (non-, weakly-, or highly metastatic) of clonal populations enriched for LOY or ROY (>80% purity).

